# The evolution of the HIV-1 integrase through constraints from folding stability and functional interactions

**DOI:** 10.64898/2026.11.16.766921

**Authors:** Valeria Zoni, Sara Cabaleiro-Martínez, David Ferreiro, Miguel Arenas

**Affiliations:** CINBIO, University of Vigo, 36310 Vigo, Spain; Department of Biochemistry, Genetics and Immunology, University of Vigo, 36310 Vigo, Spain

**Keywords:** Protein evolution, phylogenetics, molecular adaptation, protein folding stability, molecular dynamics simulations, protein interactions

## Abstract

The rapid evolution of human immunodeficiency virus type 1 (HIV-1) facilitates the emergence and fixation of drug-resistance mutations (DRMs). Although variants carrying DRMs gain a selective advantage under antiretroviral treatment, these mutations may compromise protein fitness, affecting viral replication efficiency and shaping evolutionary trajectories. Understanding the emergence, fixation, and fitness effects of DRMs is therefore essential for predicting evolutionary outcomes and optimizing therapies. Here, we investigated the evolution of the HIV-1 integrase, a key antiretroviral target, and the influence of common DRMs by analyzing folding stability, DNA substrate binding affinity, and structural properties across integrase variants using long molecular dynamics simulations combined with protein phylogenetic inference. DRMs generally altered protein folding stability and protein-DNA interactions to a marginal extent, due to their number, position, and physicochemical properties. Several DRMs altered catalytic site geometry, including alteration of magnesium coordination within the catalytic triad and modulation of enzymatic activity by changes in flexibility of the 140–149 loop involved in substrate binding, while others had minor effects on DNA binding. Over long evolutionary timescales, temporal fluctuations in folding stability and DNA-binding affinity were observed, potentially driven by alternating diversification and selection regimes at molecular and population levels. We conclude that HIV-1 integrase evolves under marginal stability, allowing structural variation that supports adaptation while preserving essential activity, which should be considered in predictions of structural evolutionary trajectories. Together, integrating biophysical properties with evolutionary modeling provides a robust framework for understanding how structural constraints influence the fixation of adaptive mutations.

**Statement:** Viral adaptation to antiretroviral therapies is a major health challenge. Understanding how HIV changes over time and develops drug resistance is essential for improving treatment strategies. Here, we studied HIV-1 integrase, an important target of antiretroviral drugs, and found that its drug-resistant variants can tolerate structural changes while maintaining their stability and function. This flexibility may help the virus adapt while preserving essential activity.

## Introduction

Infections caused by viruses resistant to antiviral treatments remain a major challenge in modern medicine (Kurt Yilmaz & Schiffer, 2021). Viral evolution can be influenced by the selective pressures imposed by therapeutic interventions, where variants carrying adaptive mutations (i.e., drug-resistance mutations, DRMs) can become fixed in the viral population (Lorenzo-Redondo et al., 2016; Moyano et al., 2022). Importantly, DRMs should not be considered solely as treatment-associated mutations, as they can arise spontaneously and are also observed in treatment-naïve patients (Parisi et al., 2007). These mutations allow viral target proteins to escape treatment inhibition while maintaining sufficient functional activity (Harrigan & Alexander, 1999), favoring their fixation in viral populations and ultimately reducing treatment efficacy and complicating disease management. However, although DRMs may confer a selective advantage under treatment, they can also compromise protein fitness (Martinez-Picado & Martínez, 2008; Menéndez-Arias et al., 2003; Quiñones-Mateu & Arts, 2002). In some cases, after the fixation of a DRM, protein fitness can subsequently be restored through compensatory mutations (González-Ortega et al., 2011; Moyano et al., 2022; Wood et al., 2013), showing the capacity of some viral proteins to mitigate consequences from treatment escape (Gong et al., 2013; Klein et al., 2018; Olabode et al., 2017). A model virus that exemplifies these challenges is the human immunodeficiency virus type 1 (HIV-1), which has a generally rapid replication cycle and whose reverse transcriptase lacks proofreading activity, leading to high mutation rates and the rapid emergence of genetic variants that facilitate adaptation to new environments (Perelson et al., 1996; Yeo et al., 2020). Current antiretroviral treatments target several HIV-1 enzymes essential for viral replication, including protease (PR), integrase (IN) and reverse transcriptase (RT) (Alan Engelman & Cherepanov, 2012; Arts & Hazuda, 2012; Iyidogan & Anderson, 2014; Maeda et al., 2019). Despite the high efficacy of these treatments, challenges such as limited drug penetration into certain reservoirs and the fixation of DRMs hinder the extinction of HIV-1 populations within hosts (Harrigan & Alexander, 1999; Lorenzo-Redondo et al., 2016). Indeed, the detection of DRMs in these proteins often indicates the need for treatment modification (Foka & Mufhandu, 2023). In this regard, strategies that anticipate evolutionarily expected variants and incorporate this information into treatment design could remain effective over longer periods. However, efforts in this direction, such as vaccines based on centralized genes (Doria-Rose et al., 2005; Nickle et al., 2003), have been only partially effective, likely due to the use of unrealistic models of HIV-1 protein evolution that rely solely on sequence information (Arenas & Posada, 2010). Substitution models of protein evolution that account for evolutionary constraints from the protein structure and activity have demonstrated more accurate evolutionary predictions (Bordner & Mittelmann, 2014; Ferreiro et al., 2024; Lorca-Alonso et al., 2025). Therefore, understanding and integrating these evolutionary constraints into substitution models of protein evolution is useful for improving the accuracy of evolutionary predictions and, by extension, the application to treatment design.

Previous studies focused on the HIV-1 PR have shown that this protein can accommodate multiple DRMs without dramatically compromising its stability, which fluctuates within a functional range (Ferreiro et al., 2022; Hamacher, 2008; Olabode et al., 2017). However, these studies ignored the effects of DRMs on protein-ligand binding interactions, which may exert selective pressure in regions near the binding site (Ferreiro et al., 2024). Consequently, both stability and binding interactions should be considered for accurately modeling viral protein evolution, with implications for the design of antiretroviral therapies (Arenas & Posada, 2010; Meixenberger et al., 2017).

In this study, we focused on the HIV-1 IN, a key enzyme responsible for integrating viral DNA into the host genome (Alan Engelman & Cherepanov, 2012). Some antiretroviral therapies target IN activity using integrase strand transfer inhibitors (INSTIs) (Espeseth et al., 2000). However, INSTI-associated DRMs have been reported (Jóźwik et al., 2020; Oliveira et al., 2018), and understanding how these mutations affect the protein fitness is important for designing effective and durable drugs. In this regard, studies investigating HIV-1 IN evolution that consider both folding stability and DNA-binding interactions remain scarce, likely due to the complexity of modeling the evolution of its interactions with the DNA substrate. Here, we examined the influence of clinically relevant INSTI resistance-associated mutations on the folding stability and DNA-binding affinity of HIV-1 IN. In addition, we explored the evolution of these properties across the long-term evolutionary history of the virus, identifying the ranges compatible with viral survival, temporal trends, and variation among viral lineages. To investigate these evolutionary constraints, we employed molecular dynamics (MD) simulations, a powerful method that captures the dynamic behavior of biomolecular structures (Frenkel & Smit, 2023) and is well suited for evaluating both molecular binding interactions and protein folding stability (Sapozhnikov et al., 2023; Wang et al., 2018), together with probabilistic methods of protein phylogenetic inference.

## Results

### Drug-resistance mutations affect the folding stability and substrate-binding affinity of HIV-1 integrase

We evaluated the folding stability and DNA substrate-binding free energy of the HIV-1 IN WT variant (Figure 1A) and of HIV-1 IN variants carrying single or combined DRMs (Figure 1B, Table 1). Overall, most variants containing one or more DRMs displayed folding stability comparable to that of the WT (Figure 2). Approximately 20% of the variants exhibited increased stability (lower ΔG), whereas only two variants showed reduced stability relative to the WT (higher ΔG) (Figure 2). Among single and combined DRMs, the largest contributions to increased stability were observed for N155H, Q148R/H/K, S147G, G140S, and R263K (Figure S1A; Supplementary Material), which also occurred most frequently either alone or in combination with other mutations. In contrast, the G140C mutation alone resulted in decreased stability, as did the E92Q+R263K combination. Notably, Q148K and Q148R restored stability when present in combination with G140C, yielding values comparable to those of the WT. Similarly, the stabilizing effect of N155H in combination with E92Q+R263K was pronounced, as the E92Q+N155H+R263K variant displayed higher stability than the WT. Several mutations, including those at positions T66 and Y143, had little or no effect on IN stability when present individually. However, Y143R exhibited a marked destabilizing effect when included in the E92Q+G140S+Q148R+N155H combination, resulting in lower stability compared with the corresponding four-mutation variant lacking Y143R.

**Figure 1.**
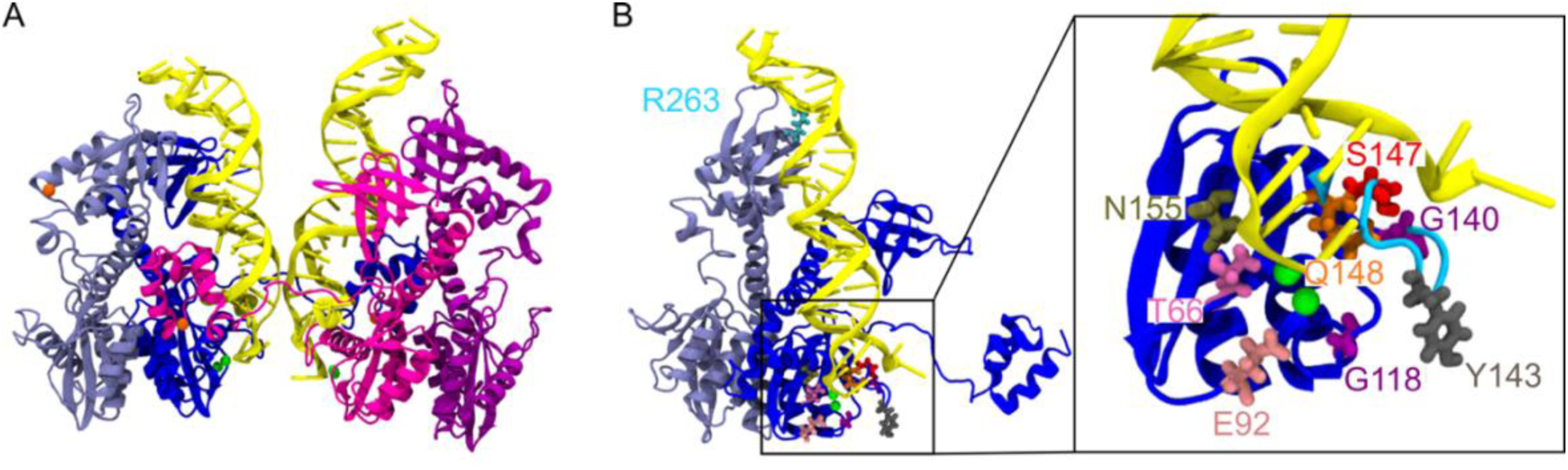
Molecular structure of the HIV-1 integrase-DNA complex and location of the analyzed drug-resistance mutations. (A) Structure of the HIV-1 integrase-DNA complex used in this study. The four integrase chains are shown in different colors: chain A in blue, chain B in blue-gray, chain C in magenta, and chain D in purple. DNA strands, Mg^2+^ ions and Zn^2+^ ions are shown in yellow, green, and orange, respectively. (B) Positions of the analyzed drug-resistance mutations in chains A and B. The WT residue R263 is highlighted in chain B and the WT residues T66, E92, G118, G140, Y143, S147, Q148 and N155 in chain A are highlighted in the inset. Chains C and D have the same residue positioning as chains A and B. The 140-149 loop is highlighted in light blue in the inset.

**Figure 2.**
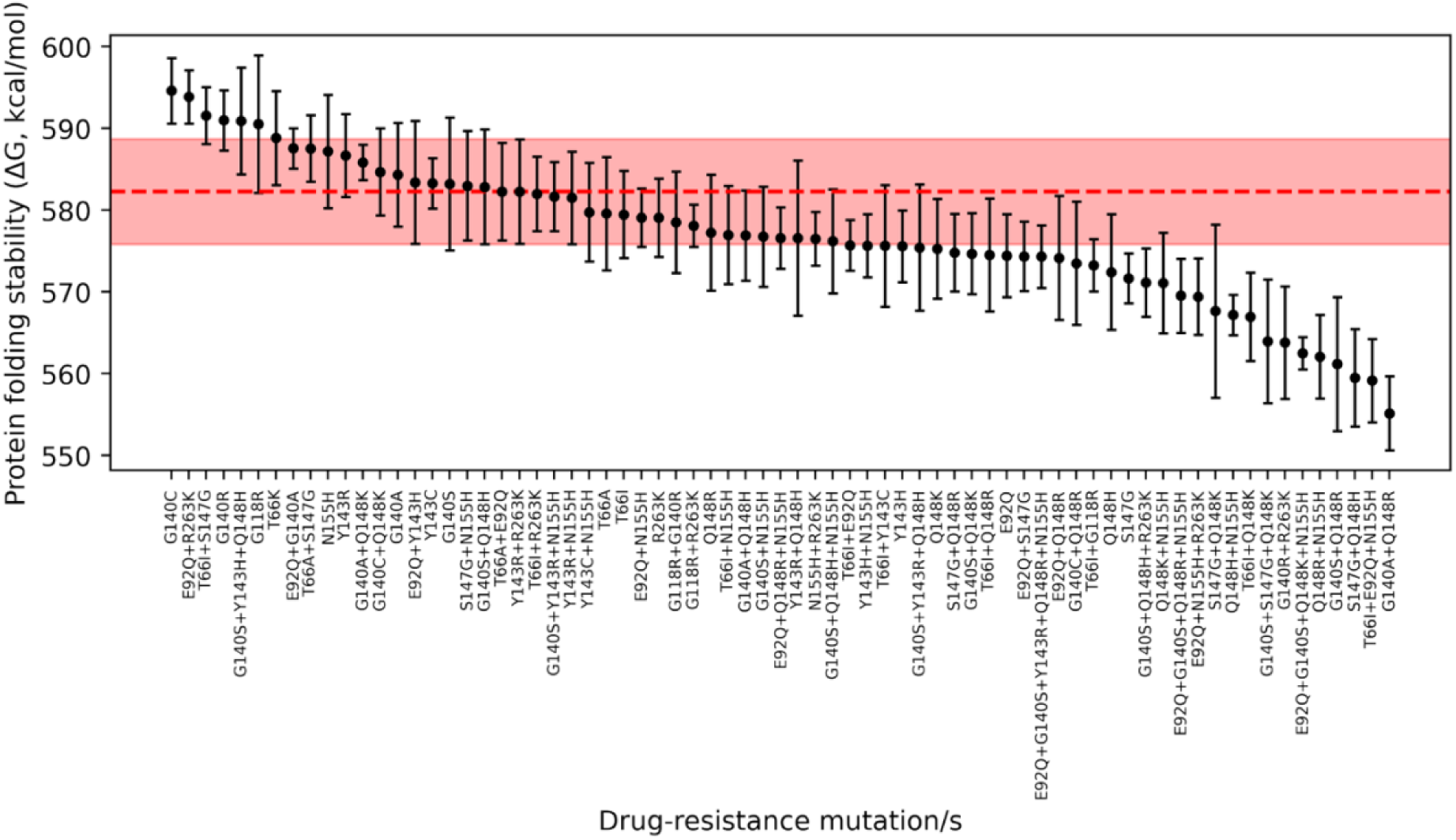
Folding stability of HIV-1 integrase variants with and without drug-resistance mutations. The plot shows the folding stability for each HIV-1 IN variant without (WT) or with one or more drug-resistance mutations. Each variant, including the WT, was analyzed in five independent replicates, and the plot displays the mean values with standard errors (error bars). For the WT variant, the mean stability is indicated by a dashed red line, and its standard error is represented by a red shaded area.

**Table 1.** Drug resistance mutations of the HIV-1 integrase analyzed in this study. Each studied drug resistance mutation (DRM) is reported together with the number of sequences it was observed in viral samples collected from patients, and the corresponding susceptibility level (scores ranging from 1 –no detected loss of susceptibility– to 5 –high-level resistance–) to different INSTIs (BIC: bictegravir; CAB: cabotegravir; DTG: dolutegravir; EVG: elvitegravir; RAL: raltegravir). An asterisk indicates combined DRMs that were observed but without a sequence available in the database. All data were retrieved from the Stanford HIV Drug Resistance Database.

|  |  | **Susceptibility level** |  |  |  |  |
|---|---|---|---|---|---|---|
| **Drug resistance mutation/s** | **Number of Sequences** | **BIC** | **CAB** | **DTG** | **EVG** | **RAL** |
| **E92Q** | 68 | 2 | 3 | 2 | 5 | 4 |
| **E92Q+G140A** | 1 | 3 | 4 | 3 | 5 | 5 |
| **E92Q+G140S+Q148K+N155H** | 3 | 5 | 5 | 5 | 5 | 5 |
| **E92Q+G140S+Q148R+N155H** | 3 | 5 | 5 | 5 | 5 | 5 |
| **E92Q+G140S+Y143R+Q148R+N155H** | 1 | 5 | 5 | 5 | 5 | 5 |
| **E92Q+N155H** | 49 | 4 | 5 | 4 | 5 | 5 |
| **E92Q+N155H+R263K** | 1 | 5 | 5 | 5 | 5 | 5 |
| **E92Q+Q148R** | 1 | 4 | 5 | 4 | 5 | 5 |
| **E92Q+Q148R+N155H** | 2 | 5 | 5 | 5 | 5 | 5 |
| **E92Q+R263K** | 1 | 4 | 5 | 4 | 5 | 4 |
| **E92Q+S147G** | 2 | 3 | 4 | 3 | 5 | 4 |
| **E92Q+Y143H** | 1 | 2 | 3 | 2 | 5 | 5 |
| **G118R** | 12 | 4 | 5 | 5 | 5 | 5 |
| **G118R+G140R** | 2 | 4 | 5 | 5 | 5 | 5 |
| **G118R+R263K** | 1 | 5 | 5 | 5 | 5 | 5 |
| **G140A** | 3 | 2 | 3 | 2 | 4 | 4 |
| **G140A+Q148H** | 2 | 4 | 5 | 4 | 5 | 5 |
| **G140A+Q148K** | 3 | 4 | 5 | 4 | 5 | 5 |
| **G140A+Q148R** | 17 | 4 | 5 | 4 | 5 | 5 |
| **G140C** | 1 | 2 | 3 | 2 | 4 | 4 |
| **G140C+Q148K\*** |  | 2 | 1 | 2 | 1 | 1 |
| **G140C+Q148R** | 8 | 4 | 5 | 4 | 5 | 5 |
| **G140R** | 47 | 2 | 5 | 2 | 4 | 4 |
| **G140R+R263K** | 1 | 4 | 5 | 4 | 5 | 4 |
| **G140S** | 16 | 2 | 3 | 2 | 4 | 4 |
| **G140S+N155H** | 2 | 3 | 4 | 3 | 5 | 5 |
| **G140S+Q148H** | 324 | 4 | 5 | 4 | 5 | 5 |
| **G140S+Q148H+N155H** | 8 | 5 | 5 | 5 | 5 | 5 |
| **G140S+Q148H+R263K** | 1 | 5 | 5 | 5 | 5 | 5 |
| **G140S+Q148K** | 6 | 4 | 5 | 4 | 5 | 5 |
| **G140S+Q148R** | 32 | 4 | 5 | 4 | 5 | 5 |
| **G140S+S147G+Q148K** | 1 | 5 | 5 | 5 | 5 | 5 |
| **G140S+Y143H+Q148H** | 5 | 4 | 5 | 4 | 5 | 5 |
| **G140S+Y143R+N155H** | 1 | 3 | 4 | 3 | 5 | 5 |
| **G140S+Y143R+Q148H** | 2 | 4 | 5 | 4 | 5 | 5 |
| **N155H** | 302 | 2 | 3 | 2 | 5 | 5 |
| **N155H+R263K** | 1 | 5 | 5 | 5 | 5 | 5 |
| **Q148H** | 32 | 3 | 5 | 3 | 5 | 5 |
| **Q148H+N155H** | 1 | 4 | 5 | 4 | 5 | 5 |
| **Q148K** | 11 | 4 | 5 | 4 | 5 | 5 |
| **Q148K+N155H\*** |  | 3 | 1 | 3 | 1 | 1 |
| **Q148R** | 52 | 3 | 5 | 3 | 5 | 5 |
| **Q148R+N155H** | 13 | 4 | 5 | 4 | 5 | 5 |
| **R263K** | 71 | 4 | 5 | 4 | 4 | 3 |
| **S147G** | 18 | 2 | 3 | 2 | 5 | 2 |
| **S147G+N155H\*** |  | 2 | 3 | 2 | 1 | 1 |
| **S147G+Q148H\*** |  | 2 | 2 | 2 | 1 | 1 |
| **S147G+Q148K\*** |  | 2 | 2 | 2 | 1 | 1 |
| **S147G+Q148R** | 1 | 4 | 5 | 4 | 5 | 5 |
| **T66A** | 18 | 1 | 1 | 1 | 5 | 3 |
| **T66A+E92Q** | 1 | 3 | 3 | 3 | 5 | 4 |
| **T66A+S147G** | 1 | 3 | 3 | 3 | 5 | 3 |
| **T66I** | 29 | 1 | 2 | 1 | 5 | 3 |
| **T66I+E92Q** | 6 | 3 | 3 | 3 | 5 | 4 |
| **T66I+E92Q+N155H** | 1 | 4 | 5 | 4 | 5 | 5 |
| **T66I+G118R** | 1 | 4 | 5 | 5 | 5 | 5 |
| **T66I+N155H** | 2 | 3 | 4 | 3 | 5 | 5 |
| **T66I+Q148K** | 1 | 4 | 5 | 4 | 5 | 5 |
| **T66I+Q148R** | 3 | 4 | 5 | 4 | 5 | 5 |
| **T66I+R263K** | 4 | 4 | 5 | 4 | 5 | 4 |
| **T66I+S147G** | 1 | 3 | 3 | 3 | 5 | 3 |
| **T66I+Y143C** | 1 | 1 | 3 | 1 | 5 | 5 |
| **T66K** | 6 | 3 | 3 | 3 | 5 | 5 |
| **Y143C** | 16 | 1 | 1 | 1 | 2 | 5 |
| **Y143C+N155H** | 2 | 2 | 4 | 2 | 5 | 5 |
| **Y143H** | 12 | 1 | 1 | 1 | 2 | 5 |
| **Y143H+N155H** | 5 | 2 | 4 | 2 | 5 | 5 |
| **Y143R** | 47 | 1 | 1 | 1 | 2 | 5 |
| **Y143R+N155H** | 4 | 2 | 4 | 2 | 5 | 5 |
| **Y143R+Q148H** | 1 | 3 | 5 | 3 | 5 | 5 |
| **Y143R+R263K** | 1 | 4 | 5 | 4 | 4 | 5 |

Despite most observed DRMs are located near the active site, where the protein interacts with the DNA substrate (Figure 1B), the overall strength of protein-substrate binding was only marginally affected by them (Figure 3). Nevertheless, a subset of DRMs significantly altered binding affinity. In particular, two variants exhibited weaker binding (higher ΔG) than the WT, whereas eleven variants showed improved binding (lower ΔG). Mutations at positions T66A/K, G140R/S/C, Q148H/K, S147G, and N155H were most frequently associated with enhanced DNA binding (Figure S1B; Supplementary Material). Notably, G140C, while destabilizing when present alone, improved binding affinity, whereas T66I had minimal impact on stability (Figure 2) but a pronounced effect on DNA binding (Figure 3).

**Figure 3.**
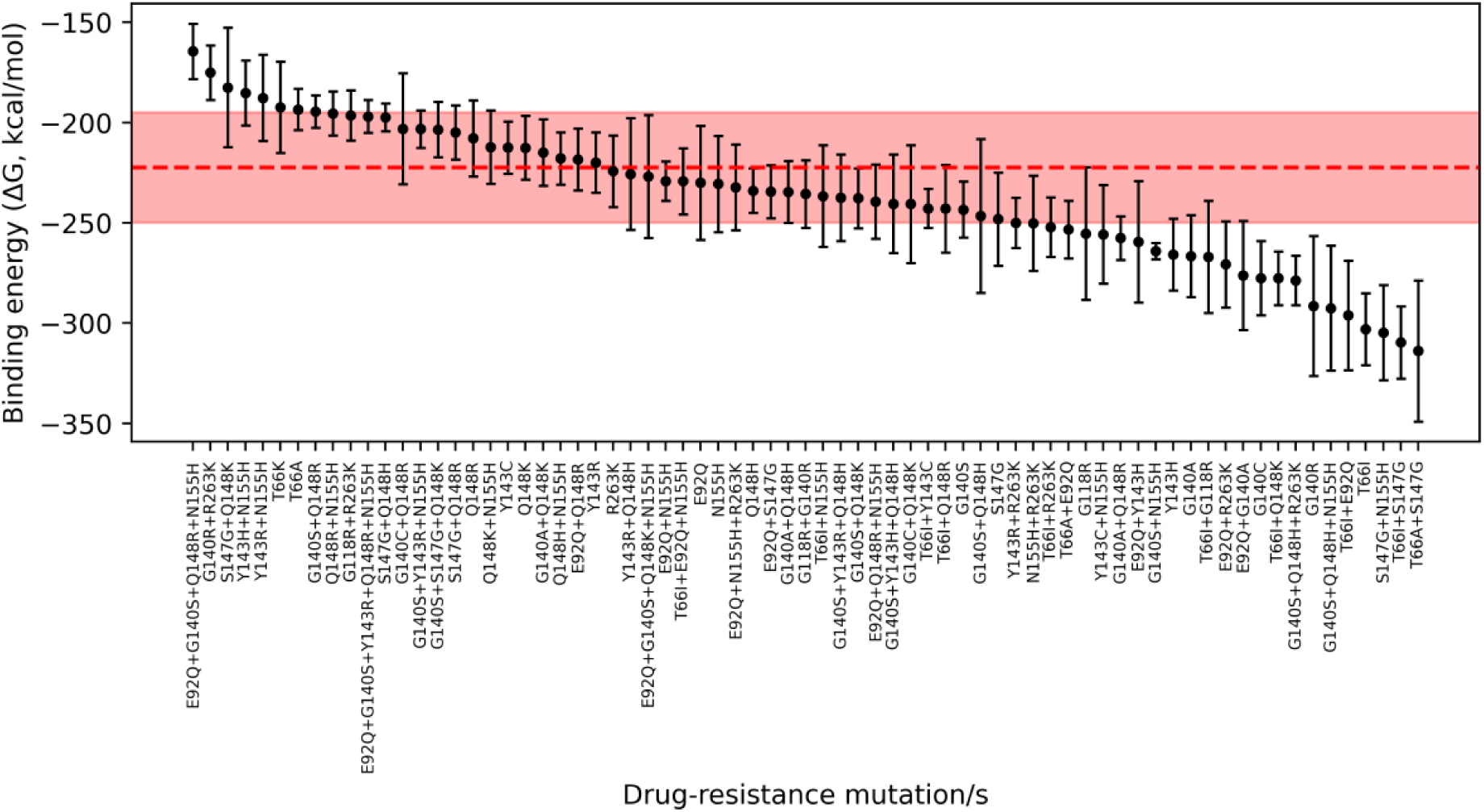
Binding free energy of HIV-1 integrase variants with and without drug-resistance mutations in complex with the DNA substrate. The plot shows the binding free energy for each HIV-1 IN variant, without (WT) or with one or more drug-resistance mutations, in complex with the DNA substrate. Each variant, including the WT, was analyzed in five independent replicates, and the mean values with standard errors (error bars) are shown. For the WT variant, the mean binding free energy is indicated by a dashed red line, and its standard error is represented by a red shaded area.

A comparison between folding stability and binding free energy revealed that variants such as G140S+Q148H+R263K and T66I+Q148K improved both properties, whereas variants including E92Q+G140S+Q148R+N155H and G140R+R263K, despite increased stability, exhibited reduced binding affinity. Finally, we investigated the contribution of epistatic effects by analyzing variants containing increasing numbers of DRMs (Figure S1C; Supplementary Material). The fraction of variants with improved stability increased progressively from one to four mutations, whereas binding affinity was more frequently improved in variants containing one to three mutations. Among four-mutation variants, one of two showed reduced binding affinity, and no clear effect was observed for the single five-mutation variant analyzed. Although the limited number of variants with four or five DRMs may bias these observations, the results suggest a role for epistasis in modulating IN stability, with combinations of two to three DRMs most frequently associated with enhanced stability.

### Drug-resistance mutations can alter the structural dynamics of the catalytic site of HIV-1 integrase

We evaluated the influence of DRMs on the structure of the catalytic site of HIV-1 IN by examining variations in the coordination between the catalytic triad and Mg^2+^ ions (Machado & Guimarães, 2020). In the WT variant, the catalytic residues form monodentate interactions with each Mg^2+^ ion, with the oxygen atoms of D64 coordinating two different Mg^2+^ ions (Figure 4, top panel). These interactions between the catalytic triad and Mg^2+^ remained stable throughout the simulations, as shown by the final frame of a representative MD simulation (Figure 4, top right) and by the persistent interactions between the oxygen atoms of D64, D116, and E152 and the Mg^2+^ ions (Figures S2 and S3; Supplementary Material). In contrast, HIV-1 IN variants carrying DRMs showed changes in the coordination of D116 and E152 with Mg^2+^ ions, while D64 interactions remained stable (Figures S2 and S3; Supplementary Material). In particular, DRMs induced spatial rearrangements of D116 and E152, as observed, for example, in variants carrying G140R and G118R (Figure 4, bottom), indicating that some mutations can alter this key structural motif and potentially affect enzymatic activity.

**Figure 4.**
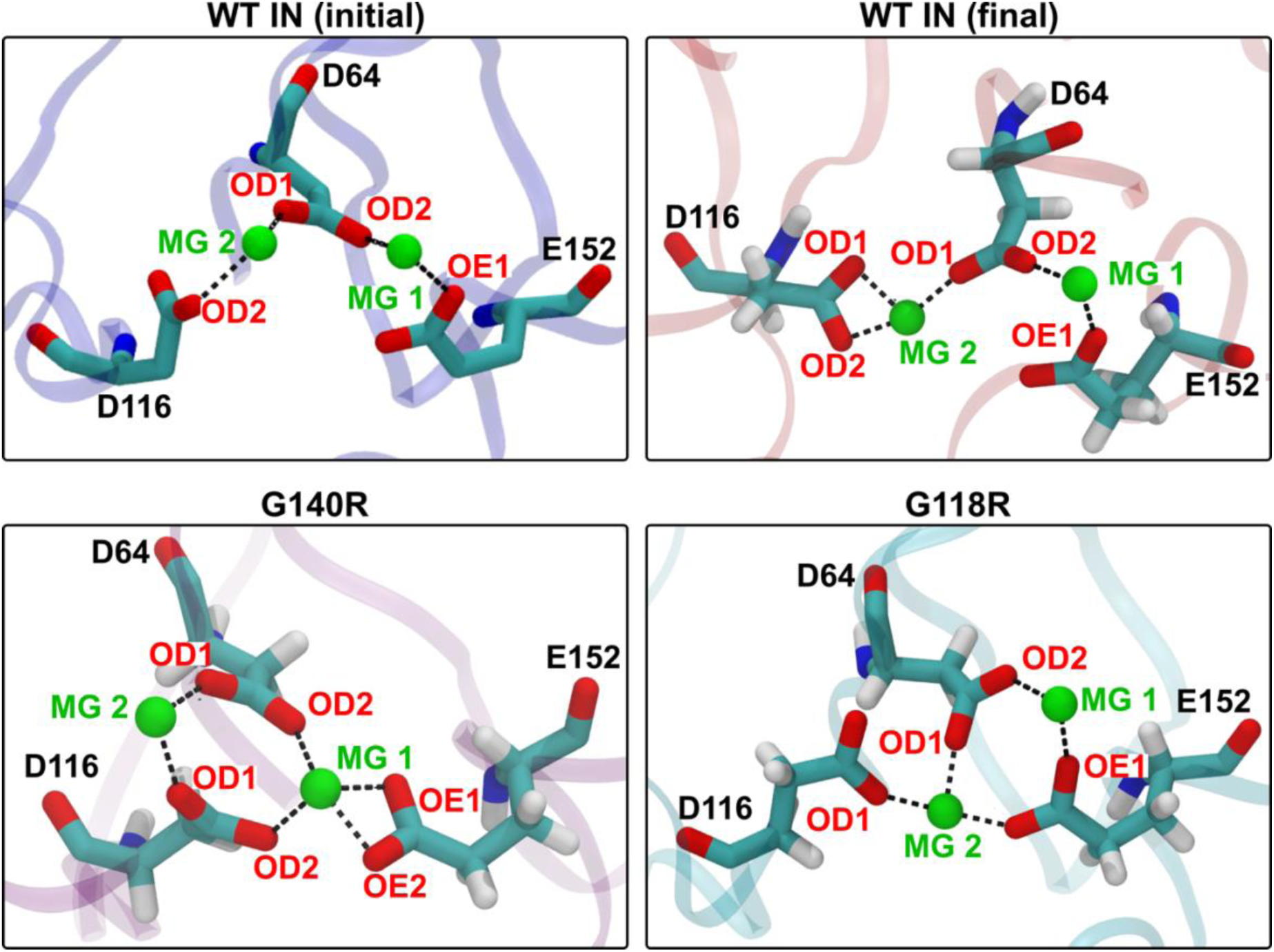
Positioning of catalytic triad and magnesium ions in HIV-1 integrase variants with and without drug-resistance mutations. The positioning of the oxygen atoms of D64 (OD1, OD2), D116 (OD1, OD2) and E152 (OE1, OE2) relative to Mg^2+^ions (MG) is shown for HIV-1 IN variants without (WT, displayed at the beginning and end of the MD simulations; top plots) or with specific drug-resistance mutations (bottom plots). For each DRM containing variant, the plot presents a representative snapshot from one replicate at the end of the MD simulation.

We also evaluated the influence of DRMs on the flexibility of the 140-149 loop (Figure 1B and Figure S4; Supplementary Material), a region implicated in DNA and drug binding (Jóźwik et al., 2020; M. Li et al., 2023; Passos et al., 2020), by calculating the RMSD of this region at the end of the MD simulations relative to the minimized structure. The results showed that HIV-1 IN variants carrying DRMs often exhibited higher RMSD variability across replicates of independent MD simulations, indicating an influence on the structural flexibility of regions involved in the protein function. In many variants with DRMs, the loop was either more flexible (higher RMSD) or more rigid (lower RMSD) than in the WT variant, additionally supporting the notion that DRMs can affect functional dynamics, which may, in turn, influence DNA or drug binding. To test this hypothesis, we calculated Pearson (Table S1; Supplementary Material) and Spearman (Table S2; Supplementary Material) correlation coefficients between folding stability or binding energy and the structural descriptors reported above (distances and RMSD). While folding stability showed little correlation with these structural indicators, we observed a notable correlation between binding energy and the distance between the OD2 oxygen of D116 and Mg²⁺ in both chains A and C. In particular, shorter D116–Mg²⁺ distances were associated with more favorable binding energies, suggesting that Mg²⁺ chelation plays a crucial role in ensuring proper binding between IN and DNA.

### Long-term evolution of the HIV-1 integrase in terms of molecular stability and binding affinity

Considering the previous effects of DRMs on the HIV-1 IN stability and binding affinity with the natural DNA substrate, we investigated the evolution of these fundamental properties along the evolutionary history of diverse HIV-1 subtypes. The results revealed multiple temporal fluctuations in both folding stability and substrate-binding affinity (Figures 5 and 6). More specifically, we observed that the evolution of folding stability varied among HIV-1 subtypes and groups, and it remained relatively constant along intermediate branches and within some subtypes (i.e., C and F of group M and group O), while being more variable within others (i.e., B) (Figure S5; Supplementary Material). Regarding the evolution of binding affinity, we also found relative preservation at intermediate branches and within some subtypes (i.e., F and D), whereas others (i.e., C of group M and group O) showed higher variability (Figure S6; Supplementary Material).

**Figure 5.**
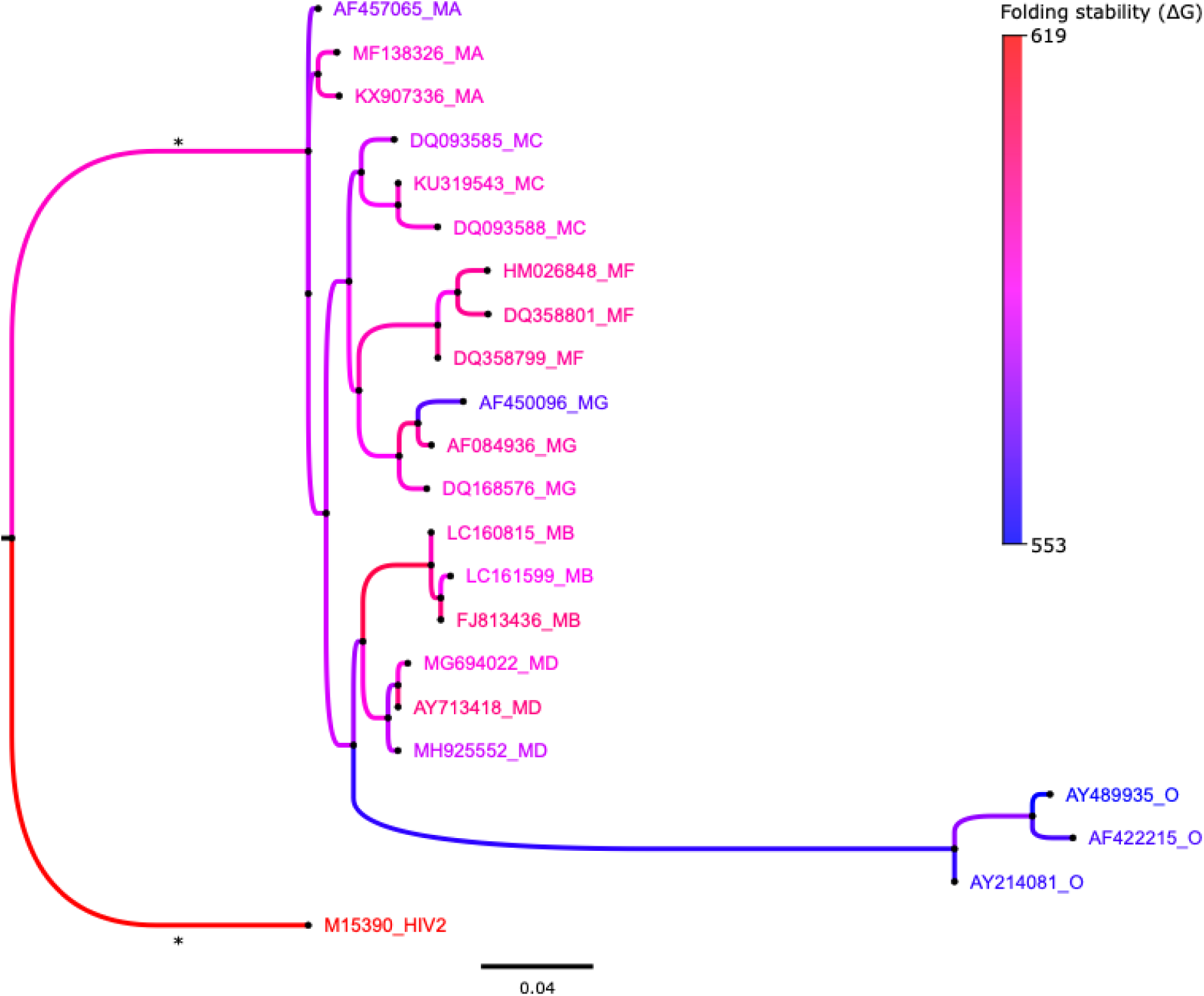
Evolution of HIV-1 integrase folding stability along the viral evolutionary history. Phylogenetic tree showing the folding stability of extant (tip nodes) and reconstructed ancestral (internal nodes) HIV-1 IN variants. Each tip node is labeled with the corresponding accession code from the HIV Drug Resistance Database (HIVdb), followed by the annotated HIV-1 group and subtype (e.g., AF457065_MA refers to HIVdb code AF457065, HIV-1 group M and, subtype A). An HIV-2 IN variant was included as an outgroup to root the tree. Phylogenetic branches are colored according to the folding stability of each variant, as indicated in the legend (note that lower ΔG values correspond to higher stability). For clarity, branches marked with an asterisk were shortened to one-fifth of their original length.

**Figure 6.**
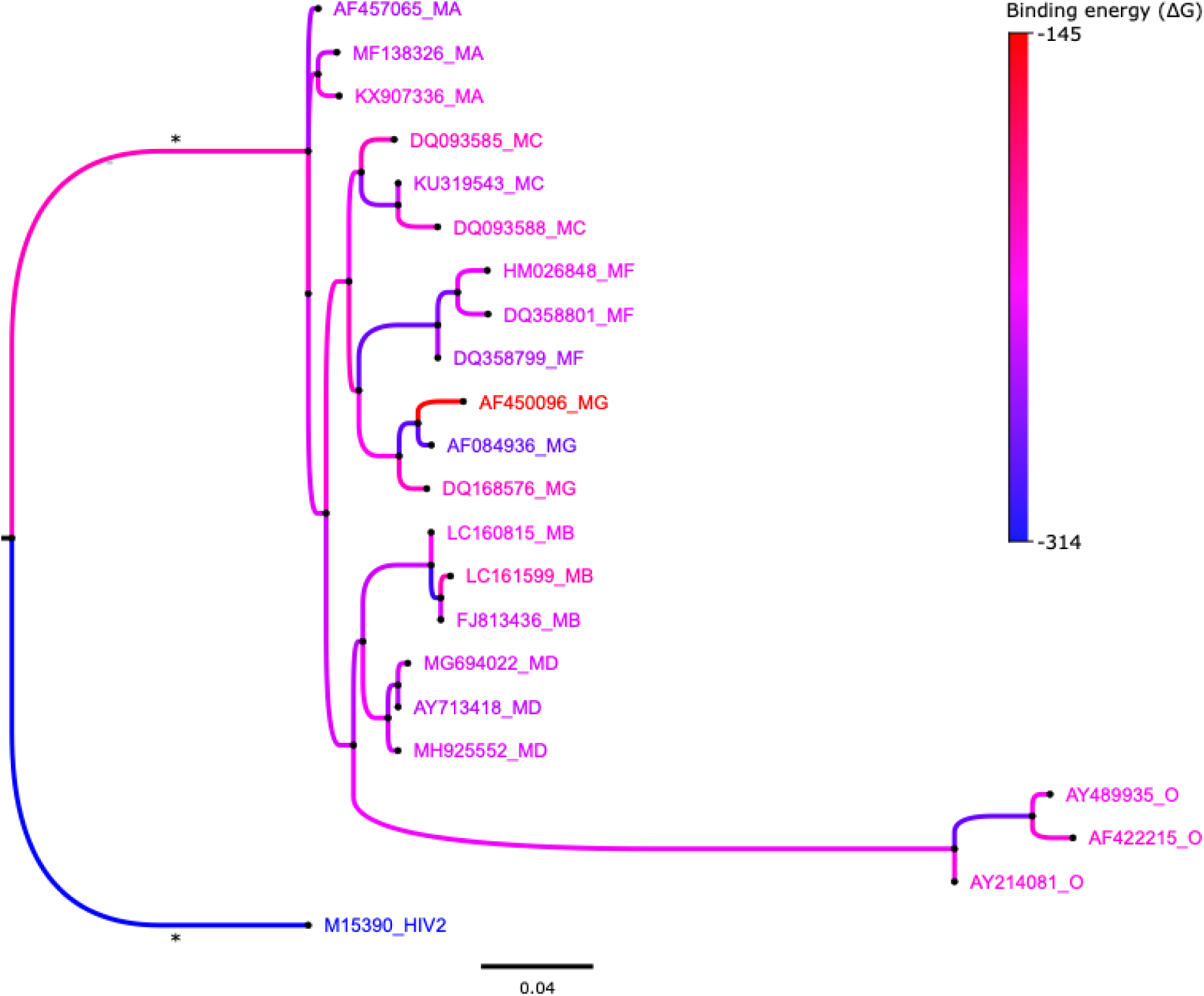
Evolution of the binding free energy of the HIV-1 integrase with its natural DNA substrate along the viral evolutionary history. Phylogenetic tree showing the binding free energy of extant (tip nodes) and reconstructed ancestral (internal nodes) HIV-1 IN variants. Each tip node is labeled with the corresponding accession code from the HIV Drug Resistance Database, followed by the annotated HIV-1 group and subtype (e.g., AF457065_MA refers to HIVdb code AF457065, HIV-1 group M and, subtype A). An HIV-2 IN variant was included as an outgroup to root the tree. Phylogenetic branches are colored according to the binding free energy (ΔG) of each variant, as indicated in the legend (note that lower ΔG values correspond to higher binding). For clarity, branches marked with an asterisk were shortened to one-fifth of their original length.

## Discussion

Understanding protein evolution enables not only the evaluation of phenotypic fitness but also the prediction of past and potential future molecular evolutionary trajectories (Ferreiro et al., 2025; Pál et al., 2006; Tokuriki & Tawfik, 2009). Indeed, viral proteins are common targets of antiretroviral therapies and, therefore, accounting for their evolutionary dynamics is essential for the design of effective treatments (Heaton, 2019; J. Zhang et al., 2010). The study of protein evolution is complex, as it is driven not only by changes in amino acid sequences but also by the consequent alterations in protein structure (Ferreiro et al., 2025; Kinch & Grishin, 2002; Pascual-García et al., 2019; Puente-Lelievre et al., 2025). In this regard, protein structures are generally more conserved than sequences (Illergård et al., 2009; Pascual-García et al., 2010), and most random mutations can destabilize the protein fold, thereby severely impairing protein function (Tokuriki et al., 2007) and typically being removed by Darwinian selection. However, many neutral mutations can still arise and persist, a situation particularly common in rapidly evolving viruses such as HIV-1. After extensive efforts and despite the challenges in developing effective global vaccines against this virus (i.e., Watkins et al., 2008), HIV-1 infection is currently managed with a wide variety of antiretroviral drugs that inhibit key viral proteins essential for replication, thereby enabling host survival (González-Navarro et al., 2024; Łupina et al., 2025). However, in addition to the known limitations of antiretroviral treatments in reaching viral reservoirs (J. Chen et al., 2022; González-Navarro et al., 2024; Kwon & Siliciano, 2017), viral proteins can accumulate mutations that allow escape not only from immune systems but also from therapies. In this regard, the specific consequences of individual and combined DRMs on the stability and activity of viral protein drug targets are not yet fully understood. Indeed, while the sequence evolution of these proteins has been widely investigated (Castro-Nallar et al., 2012; Del Amparo & Arenas, 2022b; Toccafondi et al., 2023), the evolution of their structural stability and functional activity has received less attention. This motivated us to explore the consequences of DRMs on the fitness of the HIV-1 IN, a key HIV-1 drug target, and the long-term evolution of its structural properties. In a previous study, Olabode et al. (2017) comprehensively explored the evolution of stability in some HIV protein drug targets, including IN. They found a general tendency for protein stability to decrease over time, including in resistant variants, and proposed that this trend could be driven by the widespread use of antiretroviral therapies. However, despite being an interesting study, it presented some methodological limitations. First, the ancestral protein sequences were reconstructed using an empirical substitution model of protein evolution based on nuclear globular proteins, and it is known that substitution models with poor fit to the data produce less accurate ancestral sequences (Del Amparo & Arenas, 2022a) and that empirical substitution models of protein evolution tend to generate proteins with unrealistic folding stabilities (Arenas et al., 2017; Arenas & Bastolla, 2019; Iglesias-Rivas et al., 2025). Here, we used an SCS model, which is known that provides more accurate ancestral sequences in terms of folding stability (Arenas et al., 2017; Arenas & Bastolla, 2019). Second, that study predicted protein stability without relaxing protein conformations (e.g., through energy minimization or MD simulations), which could affect estimates of folding stability (Sapozhnikov et al., 2023), and did not account for prediction errors or include statistical evaluations of the results, all aspects included in the present study. Finally, protein activity, a key evolutionary constraint due to its direct effects on protein fitness (Echave, 2019; Ferreiro et al., 2024; Ribeiro et al., 2020), was not considered in that study, while it is explicitly evaluated in the present study.

Here, we investigated the consequences of a large number of clinically crucial DRMs and their combinations (Table 1) on the stability and activity (evaluated through binding affinity with the natural substrate), as this information could help design therapies that remain effective for longer periods (Métifiot et al., 2013; Trivedi et al., 2020; Zhao et al., 2022). In addition, we examined the evolution of these fundamental properties along the evolutionary history of HIV-1 IN. Using advanced approaches for predicting molecular stability and enzyme-substrate interactions, together with accurate SCS models of protein evolution, we found that the molecular stability of the HIV-1 IN can be affected by DRMs in contrasting ways. While some DRMs preserve protein stability, others increase or decrease it. Overall, variants carrying a single DRM maintained folding stability comparable to that of the WT (Figure 2 and Figure S1C; Supplementary Material). In contrast, combinations of DRMs tended to favor stability, especially those involving Q148H/K/R together with N155H, S147G, and/or G140A/S/R (Figure S1C; Supplementary Material), suggesting that cooperative effects between DRMs play a major role. Moreover, both the position and the physicochemical nature of the substituted residues appear to be important.

To investigate the structural basis of these effects, we estimated multiple energy terms (i.e., van der Waals, electrostatic, hydrophobic solvation, and polar solvation) contributing to the total folding energy of the analyzed variants (Figure S7; Supplementary Material). When considering only the variants that exhibit increased stability relative to the WT, the results showed that the main contributions arise from improved amino acid packing, reflected by more negative van der Waals (vdW) energies and increased hydrophobic burial. This trend is particularly evident in certain substitutions at positions 140 and 148 that led to increased stability, consistent with the fact that these residues lie at opposite ends of the 140–149 loop and interact with each other (Figure 1B). In particular, stabilization was observed when a small residue at position 140 (G140A or G140S) accompanied the introduction of arginine at position 148, suggesting that reduced steric constraints at position 140 allow more favorable packing of the bulky charged side chain at 148R. Consistently, other variants combining G140S, Q148H/K and additional DRMs (E92Q+G140S+Q148K/R+N155H, G140S+S147G+Q148K, and G140S+Q148H+R263K) also showed improved stability. By contrast, S147G appeared to act as a permissive mutation that generally relieves local structural constraints, enabling multiple substitutions at position 148 to be accommodated without loss of stability. Conversely, among variants displaying reduced stability relative to the WT, the G140C substitution was associated with a strong increase in vdW clash energy, together with less favorable hydrophobic solvation and weakened electrostatic interactions (Figure S7; Supplementary Material). This is consistent with the loss of glycine-specific backbone flexibility, the introduction of a bulkier side chain and the unfavorable burial of a cysteine thiol group. Another destabilizing combination observed was E92Q+R263K, which shows pronounced steric and solvation penalties (increased vdW clashes and a strong hydrophobic penalty, Figure S7; Supplementary Material). Notably, the triple mutant E92Q+N155H+R263K exhibited increased stability compared to the WT, indicating that the presence of N155H, despite being distant from R263K, can compensate for unfavorable energetic contributions and improve the overall stability profile. Notably, the N155H+R263K mutations typically precede the acquisition of the E92Q mutation (Anstett et al., 2015).

Experimental evidence indicates that the resulting triple mutant presents both reduced infectivity and decreased drug susceptibility compared to the double mutant. While these findings may initially appear to contrast with our results since the triple mutant presents a higher stability than the WT (Figure 2), they instead suggest that the reduced viral activity can be mediated by alternative mechanisms such as altered interactions with the Mg²⁺ ions (Figures S2 and S3; Supplementary Material). Indeed, for a mutation associated with a fitness cost to be maintained in a viral population, this disadvantage could be compensated by another factor such as increased drug resistance. The examples above suggest that the effects of DRMs on HIV-1 IN are diverse and not straightforward, as they depend on the specific molecular interactions involved. As Zuckerkandl and Pauling (1965) indicated over 60 years ago, ‘*it is the type rather than number of amino acid substitutions that is decisive*’ and, together with the complex network of derived molecular interactions, we find that this applies to HIV-1 IN DRMs.

Next, since INSTIs bind at the interface between the viral DNA and the catalytic core of HIV-1 IN, interacting with residues directly involved in DNA recognition (Alan Engelman & Cherepanov, 2012; Dicker et al., 2008; Jóźwik et al., 2020; Passos et al., 2020; Smith et al., 2021), many DRMs arise in or near these regions (Passos et al., 2020), potentially impairing the interaction between the protein and its natural DNA substrate. However, the studied HIV-1 IN variants carrying DRMs were fixed in viral populations, indicating that protein activity remained at least partially functional. Accordingly, we found that most DRMs did not significantly impair the binding free energy between the protein and its natural substrate.

Still, we observed some changes in binding energy, with around 20% of the studied variants showing increased binding (Figure 3 and Figure S1B; Supplementary Material). A notable example is Q148, whose interaction with DNA has been identified as crucial for proper strand-transfer activity through direct contacts with the substrate (Johnson et al., 2006). We also found that combinations containing Q148K or Q148H generally showed increased binding to DNA, whereas Q148R had either no effect or appeared deleterious. This was particularly evident in the combination E92Q+G140S+Q148R+N155H, which exhibited reduced binding compared to the corresponding variant carrying Q148H. This effect is likely driven by steric constraints, as arginine is bulkier and may perturb optimal DNA contacts.

Consistently, analysis of the energy components contributing to the binding energy revealed increased vdW penalties in the Q148R variant relative to the Q148H counterpart (Figure S8; Supplementary Material). Analysis of other mutations showed that the G140R+R263K combination was stabilizing but displayed weaker binding than WT, probably caused by R263K that was suggested to modify the binding to DNA *in vitro* (Quashie et al., 2012). In this case, the reduced binding affinity is mainly driven by increased polar solvation penalties that outweigh strengthened electrostatic interactions, while modest changes in vdW and nonpolar solvation contributions were insufficient to compensate (Figure S8; Supplementary Material). Finally, two combinations appeared to improve both binding and stability, G140S+Q148H+R263K and T66I+Q148K. For G140S+Q148H+R263K, improved packing contributes to increased binding, reflected by more favorable vdW energies, together with an overall stabilization of the folded state (improved vdW and electrostatics, increased hydrophobic burial, and reduced steric strain, Figures S7 and S8; Supplementary Material). In the case of T66I+Q148K, mutations at position 66 generally did not strongly affect stability. However, T66I provided a modest stabilizing contribution by improving vdW packing and nonpolar surface burial. This effect appears sufficient to partially compensate for the unfavorable contribution of Q148K alone, which is associated with destabilizing electrostatic effects and solvation penalties.

Interestingly, we found that certain DRM combinations improved both stability and binding affinity, consistent with experimental findings showing that DNA binding by IN variants carrying DRMs can remain preserved (Delelis et al., 2010; Delelis, Malet, Na, et al., 2009; Dicker et al., 2008; Engelman & Craigie, 1992). However, some DRMs are known to affect the catalytic activity of HIV-1 IN (Abram et al., 2013; Dicker et al., 2008; M. Li et al., 2023), which is mediated by the catalytic triad (D64, D116, and E152) via interactions with Mg^2+^ ions and DNA (Engelman & Craigie, 1992; Goldgur et al., 1998). Therefore, we analyzed two key structural features linked to the HIV-1 IN activity: (*i*) the spatial arrangement of the catalytic triad and the essential Mg^2+^ ions, which could be disrupted by certain DRMs (Machado & Guimarães, 2020), and (*ii*) the flexibility of the 140–149 loop, which is involved in both natural substrate and drug binding (Q. Chen et al., 2013; Greenwald et al., 1999; Perryman et al., 2010). Our results indicated that DRMs do not directly affect the catalytic triad but can significantly alter protein structural dynamics and the conformation of the active site. In particular, deviations in the interactions between the catalytic triad and Mg^2+^ ions observed in several variants carrying DRMs may impair catalytic activity without disrupting DNA binding, which is consistent with experimental studies reporting decreased enzymatic activity despite intact DNA affinity (Delelis, Malet, Na, et al., 2009; Delelis et al., 2010; Dicker et al., 2008). Notably, we observed a correlation between the distance between OD2 oxygen of D116 and Mg^2+^ in both chains A and C, suggesting that the positioning of Mg^2+^ ions is important for proper DNA binding. For example, several studies (Delelis, Malet, Na, et al., 2009; Delelis, Malet, Tchertanov, et al., 2009; Hu & Kuritzkes, 2010; Shadrina et al., 2015) have shown that the catalytic impairment caused by Q148H or Q148K is rescued by G140S (Q148H+G140S, Q148K+G140S), and that this impairment is not observed when G140S, E92Q, or N155H are assessed individually. Our measurements of this distance indicated that only the Q148H and Q148K single mutants differ significantly from WT, whereas the distance is restored in the Q148H+G140S and Q148K+G140S double mutants (Figures S2 and S3; Supplementary Material). However, it should be noted that another study reported lower integration activity for the G140S+Q148H and N155H variants (Varadarajan et al., 2016). Our results are consistent with observations from the same study for Y143R, for which no significant differences in the positioning of the Mg²⁺ ions were observed (Varadarajan et al., 2016). Finally, studies of G118R have reported a marked reduction in IN activity, and our simulations revealed a pronounced deviation of the OD2 atom of D116 and the Mg²⁺ ion in chain C, which is consistent with these experimental findings (Shadrina et al., 2015). These conformational changes in the catalytic site might also explain why certain protein variants are resistant to drugs, as INSTIs bind near this site and interact directly with Mg^2+^ ions (Machado & Guimarães, 2020; Passos et al., 2020). Additionally, several DRMs increased the structural variability of the 140–149 loop, as quantified by RMSD (Figure S4; Supplementary Material). Overall, these results indicate that DRMs can alter the spatial organization of key residues within the catalytic region, without directly impairing DNA binding, but affecting the rigidity of the 140–149 loop or promoting an unfavorable alignment of the catalytic triad relative to the Mg^2+^ ions, which may hinder the enzymatic activity without directly impairing DNA binding.

During the evolutionary trajectories of the HIV-1 IN, viral populations frequently overcome deleterious effects of DRMs through the fixation of compensatory mutations that restore structural integrity and catalytic activity (Ferreiro et al., 2022, 2024). Here, we detected the synergistic interaction between the G140S and Q148R substitutions, which not only increases the stability and binding of the complex but also reduces the susceptibility to INSTIs (Cook et al., 2020; D’Antoni et al., 2025; M. Li et al., 2023; W. W. Zhang et al., 2018). Indeed, Li et al. (2023) proved coevolution of G140S/Q148H and G140A/Q148K substitutions, which may indicate that protein stability is a functional constraint along the evolutionary history of HIV IN.

We also explored the long-term evolution of the HIV-1 IN stability and DNA-binding affinity by analyzing extant and reconstructed ancestral variants from different HIV-1 subgroups and subtypes. Interestingly, by considering real-world sequences derived from HIVdb (Shafer, 2006), we found that the study variants describe functional and viable evolutionary pathways rather than transient intermediates. Since the reconstruction of ancestral protein sequences is influenced by selection from the protein structure (i.e., Arenas et al., 2015; Arenas & Bastolla, 2019), integrating these biophysical constraints is required to provide robust information on the HIV-1 IN evolution, extending previous studies based on less realistic substitution models and other proteins (Ferreiro et al., 2022, 2024; Olabode et al., 2017). We found temporal fluctuations in folding stability and DNA-binding affinity both within and among subtypes, but these remained within a consistent range, suggesting that HIV-1 IN is tolerant to accommodate mutations as long as the resulting stability remained within viable levels, which is consistent with marginal stability (Goldstein, 2011; Martin & Vila, 2020; Taverna & Goldstein, 2002). This general maintenance of HIV IN stability contrasts with the findings of Olabode et al., (2017) that indicated a general decrease in stability over time. As discussed above, this discrepancy may result from their use of an empirical substitution model not based on HIV data [i.e., WAG (Whelan & Goldman, 2001)] and that assumes a single exchangeability matrix for all protein sites. It is known that model misspecification can reduce the accuracy of ancestral sequence reconstruction (Arenas et al., 2017; Arenas & Bastolla, 2019; Del Amparo & Arenas, 2022a). We employed a substitution model that incorporates evolutionary constraints from the HIV-1 IN structure and generates site-specific exchangeability matrices, which have been found to outperform empirical models in previous studies (Arenas et al., 2015; Arenas & Bastolla, 2019; Ferreiro et al., 2026). Furthermore, by applying MD simulations, we predicted folding stability while accounting for multiple structural conformations capturing mutational-induced structural shifts that remain inaccessible to simple energy minimization techniques (Sapozhnikov et al., 2023). Over long evolutionary timescales, additional lineage-specific biological processes could also promote variations in protein stability across diverse proteins, as discussed in previous studies (Arenas et al., 2017; Chi & Liberles, 2016). These include changes in effective population size that influence the strength of natural selection, variation in environmental conditions such as temperature (Serohijos & Shakhnovich, 2014), some shifts in the mutation rate and mutation patterns (Mendez et al., 2010), and episodes of diversifying selection driven by specific functional properties (Pascual-García et al., 2010).

Altogether, HIV IN stability and activity vary among a wide variety of DRM-carrying variants and over time, exhibiting both stabilizing and destabilizing effects within a marginal range likely constrained by the minimum levels of protein stability and activity required for viral survival. Variants lacking sufficient stability or activity would not be observed due to natural selection. DRM effects depended on their specific number, position, and physicochemical properties, with some DRMs even altering catalytic site geometry and modulating enzymatic activity, while others had minor effects on folding stability and DNA binding. These findings indicate that forecasting HIV IN DRMs likely to arise over time within a viral population, a promising strategy currently explored that could guide therapy selection (Aggarwal & Periwal, 2026; Huot et al., 2025; Shahin et al., 2026), is not straightforward since the fitness landscape based on these fundamental structural properties permits numerous diverse DRM variants. Moreover, our results indicate that structural flexibility (Figures S2.1 to S2.12, Supplementary Material 2)can be accommodated without compromising the stability or function of the protein, consistent with the concept of marginal stability, whereby proteins maintain a balance between structural plasticity and the stability required for biological function (Taverna & Goldstein, 2002; Williams et al., 2007). The preservation of protein stability and DNA binding despite the structural variations observed across molecular dynamics simulations of drug-resistance, patient-derived and ancestral variants supports the presence of evolutionary constraints that maintain the functional properties of HIV-1 integrase. Finally, we emphasize the importance of incorporating dynamic structural information into evolutionary and functional analyses (Sapozhnikov et al., 2023). Unlike static structure-based predictions, MD simulations capture the flexibility and conformational rearrangements of key functional regions, including regions harboring DRMs. Although computationally demanding, MD-based approaches are essential for assessing the subtle yet functionally relevant effects that mutations can have on complex molecular assemblies such as the HIV-1 IN-DNA interface.

## Materials and methods

### General setup of HIV-1 integrase structures

We investigated the effects of DRMs on HIV-1 IN using all-atom MD simulations of the intasome, a nucleoprotein complex targeted by INSTIs (Espeseth et al., 2000). The intasome comprises 4 IN subunits arranged as 2 symmetric pairs of dimers, 2 strands of viral DNA, 4 catalytic magnesium ions (Mg^2+^), and 4 zinc ions (Zn^2+^) coordinated within the zinc-binding domain (Figure 1) (Alan Engelman & Cherepanov, 2012). We used as a reference a sequence without DRMs (hereafter referred to as wild type, WT) obtained from the Stanford HIV Drug Resistance Database (HIVdb) (Shafer, 2006). In order to choose the appropriate structure for our studies, we took into account that the assembly of the intasome and the catalytic activity of IN follow a multi-step process (Passos et al., 2020). We chose structures of the cleaved synaptic complex, which is catalytically active and the specific target of INSTIs, that were obtained in the absence of drugs to avoid potential drug-induced distortions of the secondary structure and its interactions with the DNA substrate (Espeseth et al., 2000; Hare et al., 2010; Xue et al., 2012). Since no entire HIV-1 IN structures covering the entire protein sequence of interest (from residue 1 to 270, excluding the C-terminal loop) are available in databases, we assembled the structure by combining the PDB structures 6PUT (Passos et al., 2020) covering residues 1 to 226, and 6VDK (M. Li et al., 2020) covering residues 227 to 270. Missing loops (residues 49 to 56 and 140 to 149 in chains B and D) were first modeled using *CHARMM-GUI* (Jo et al., 2008, 2014) and subsequently refined with *Modeller* (Fiser et al., 2000; Webb & Sali, 2016). The structure and relative positioning of the DNA molecule, as well as Zn^2+^ and Mg^2+^ ions, were obtained from the cited protein structures.

Drug-resistant HIV-1 IN variants (Table 1) observed in patients were retrieved from the HIVdb and included single and combined DRMs associated with reduced susceptibility or virologic response to INSTIs, according to the database. We generated the structures of the resistant variants using *AmberTools23* (Case et al., 2023), specifically using the preparatory tools *pdb4amber* to mutate backbone atoms and *Leap* to reconstruct side chains. Additionally, intasome structures from observed and reconstructed ancestral HIV-1 IN sequences (details provided later) were obtained through homology modeling with *Modeller* (Fiser et al., 2000; Webb & Sali, 2016), selecting the structural model with the best DOPE score (lowest energy) (Shen & Sali, 2006).

### Molecular dynamics simulations of HIV-1 integrase variants

We used MD simulations, which allow the evaluation of conformational flexibility and energy fluctuations in protein structures (Adcock & McCammon, 2006; Fan & Mark, 2004; Patodia, 2014), to assess the folding stability and DNA binding affinity of HIV-1 IN variants, including those carrying DRMs and reconstructed ancestral variants. To prepare the systems for MD simulations, all HIV-1 IN structures were placed into a cuboid box with a minimum distance of 1.5 nm from any atom of the intasome and solvated with water, under the 4-point, three-charge rigid water model (OPC) (Izadi et al., 2014), and Na^+^ counterions and NaCl were added to a final concentration of 0.15 M using the *Leap* preprocessing program. We performed the MD simulations using *AMBER20* (Case et al., 2020) with the ff19SB force field for proteins (Tian et al., 2020), the OL21 force field for DNA (Zgarbová et al., 2021), and the OPC model for water (Izadi et al., 2014). Ion parameters compatible with the OPC water model were used, including monovalent ions (Sengupta et al., 2021) and divalent ions (Z. Li et al., 2020). Each system was subjected to three consecutive energy minimizations, gradually applying and then removing harmonic restraints on different molecules of the systems to achieve progressive relaxation. Each minimization consisted of 1,250 steps with the steepest descent method and 1,250 steps with the conjugated gradient method (Becker et al., 2001). Harmonic restraints were applied to the entire intasome during the first minimization, to the non-hydrogen atoms of the protein and DNA during the second, and to the protein backbone atoms (α-carbons and nitrogens), phosphate groups, Mg^2+^ and Zn^2+^ ions during the third. These energy minimizations were followed by an additional minimization, without harmonic restraints, consisting of 5,000 steps with the steepest descent method and 5,000 steps with the conjugate gradient method (Ferreiro et al., 2024). Next, the system was equilibrated for 100 ps in the canonical ensemble, with the temperature gradually increased from 0 to 310K over the first 30 ps and maintained for the remaining 70 ps (Becker et al., 2001; Braun et al., 2019; Ferreiro et al., 2024). Lastly, we performed a production of 120 ns in the isothermal-isobaric ensemble, at 1 bar pressure, controlled by the Berendsen barostat (Berendsen et al., 1984) using isotropic pressure coupling. For both the equilibration and production phases, a time step of 2 fs was used, employing Langevin dynamics with a collision frequency of 2 ps^-1^, the SHAKE algorithm (Ryckaert et al., 1977) to constrain hydrogen bond lengths, and a cutoff of 10 Å for non-bonded interactions. For each system, we performed five independent analyses, and the presented results correspond to their mean and standard error. To assess structural variations throughout the simulations, the root mean square deviation (RMSD; Figures S2.1 to S2.6, Supplementary material 2) and root mean square fluctuation (RMSF; Figures S2.7 to S2.12, Supplementary Material 2) were calculated over the entire production trajectory.

### Predictions of folding stability and binding free energy of HIV-1 integrase structural variants

We estimated the folding stability of the HIV-1 IN variants and their binding free energies with the DNA substrate accounting for interactions with Mg²⁺. In particular, the folding stability was estimated using *FoldX 5.0* (Delgado et al., 2019; Guerois et al., 2002) on the indicated last 20 ns of production. There, protein structures were extracted at 100 ps intervals (i.e., 200 structures) and folding stability was calculated for each structure. The reported total and individual component energies correspond to the average over all sampled conformations. The binding free energy between the HIV-1 IN and its natural ligands (DNA and Mg^2+^) for all structural variants obtained from the MD simulations was calculated using the Molecular Mechanics/Generalized Born surface area (MMGBSA) method (Miller et al., 2012) over the last 20 ns of production. This postprocessing approach allows calculation of the free energy for the entire complex (protein–DNA–Mg^2+^) or for its individual components, the protein and the ligand. Note that the binding free energy can be defined as the difference between the free energy of the full complex and that of the separated components.

### Assessment of catalytic triad alteration and structural divergence among HIV-1 integrase variants

We evaluated the integrity of the catalytic triad (residues D64, D116, and E152 in chains A and C) by measuring the distances between Mg^2+^ ions and the carboxylate oxygens of these residues using the CPPTRAJ module (Roe & Cheatham, 2013). Alteration was defined as any increase in average distance relative to the WT configuration. Additionally, we calculated the root mean square deviation (RMSD) using CPPTRAJ, focusing only on the α-carbon and nitrogen atoms of the backbone of the 140–149 loop in chains A and C. The reference structure for RMSD calculations was the configuration at the last step of energy minimization. All analyses were performed on the last 20 ns of the production trajectory and exclusively on chains A and C, as they constitute the chains in direct contact with the DNA ligand (Figure 1). Calculation of Pearson and Spearman correlation coefficients and p-values was performed using the Python SciPy library (Virtanen et al., 2020).

### Reconstruction of ancestral HIV-1 integrase sequences and assessment of their stability and substrate-binding affinity

We explored the long-term evolution of the HIV-1 IN in terms of stability and DNA-binding affinity using protein data available from the HIVdb. In particular, we analyzed a dataset of 21 HIV-1 IN sequences representative of diverse HIV-1 subtypes (A, B, C, D, F, and G from group M) and group O, along with an HIV-2 IN sequence used as an outgroup. Next, we identified the substitution model of protein evolution that best fit the study dataset under the Bayesian Information Criterion (BIC) (Luo et al., 2010) using *ModelTest-NG* (Darriba et al., 2020). The best-fitting model was HIVb (Nickle et al., 2007) with substitution rate variation among sites modeled by a gamma distribution and including a proportion of invariable sites (HIVb +G). We applied this model to infer a maximum-likelihood (ML) phylogenetic tree, rooted with the HIV-2 IN sequence, using *RAxML-NG* (Kozlov et al., 2019). Next, we reconstructed the ancestral protein sequences corresponding to the internal nodes of the phylogeny using a structurally constrained substitution (SCS) model (Arenas et al., 2015), as implemented in the *ProtASR2* framework (Arenas & Bastolla, 2019). This approach incorporates selection on protein folding stability based on a representative protein structure (i.e., the structure of the WT sequence). The thermodynamic parameters of the SCS model were set to their default values following the recommendations of the developers (Arenas & Bastolla, 2019). Finally, we predicted the substrate-binding affinity and folding stability of each inferred ancestral HIV-1 IN sequence using MD simulations, following the procedures described in the previous sections (including five independent replicates per studied protein sequence).

### Image rendering

All snapshots from MD simulations were rendered using the *VMD* framework (Humphrey et al., 1996).

## Abbreviations

DRMs: drug-resistance mutations
HIV-1/HIV-2: human immunodeficiency virus type 1/type 2
IN: integrase
INSTIs: integrase strand transfer inhibitors
MD: molecular dynamics
ML: maximum-likelihood
MM/GBSA: Molecular Mechanics/Generalized Born surface area
PR: protease
RMSD: root mean square deviation
RMSF: root mean square fluctuation
RT: reverse transcriptase
SCS: structurally constrained substitution
VdW: van der Waals
WT: wild type

## Supplementary Material

The supplementary materials 1 and 2 are available at the journal online. Supplementary Material 1 includes the Tables S1-S2 and Figures S1-S8, while Supplementary material 1 includes the Figures S2.1-S2.12 with analysis from molecular dynamics simulations.

## Data Availability

Additional material has been uploaded to the repository https://doi.org/10.5281/zenodo.18592321. This collection includes scripts and input files for performing the molecular dynamics (MD) simulations for all DRM variants, multiple sequence alignments, phylogenetic history, and sequences of both extant and reconstructed variants, together with the corresponding MD input files, as well as a table compiling all the results.

## Acknowledgments

This study was supported by the Grant JDC2022-048873-I funded by MICIU/AEI/10.13039/501100011033 and by the “European Union NextGenerationEU/PRTR” and, by the Grant “European Union’s Horizon Europe research and innovation programme under the Marie Skłodowska-Curie grant agreement No 101149811 (Project: PRERES)”. It was also supported by the Project PID2023-151032NB-C22 funded by MICIU/AEI/10.13039/501100011033 and by FEDER, UE. Funding for open access charge: Universidade de Vigo/CISUG. We thank the “Centro de Supercomputación de Galicia (CESGA)” for the computer resources. The funders had no role in study design, data collection and analysis, decision to publish, or preparation of the manuscript.

