## Supplementary Information 1 for "The evolution of the HIV-1 integrase through constraints from folding stability and functional interactions"

The supplementary material includes the Tables S1-S2 and Figures S1-S8.

### Supplementary Tables

**Table S1. Pearson correlation coefficients between folding stability or binding energy and structural descriptors.** Structural descriptors include distances between the oxygen atoms of catalytic triad and $Mg^{2+}$ ions, as well as the RMSD of the 140–149 loop. Corresponding *p*-values are reported in parentheses, and statistically significant correlations (p-value < 0.01) are highlighted in red.

| Structural descriptor | Pearson correlation with folding stability (p-value) | Pearson correlation with binding energy (p-value) |
|---|---|---|
| **RMSD chain A** | 0.251 ($3.371\times10^{-2}$) | -0.067 ($5.789\times10^{-1}$) |
| **RMSD chain C** | 0.104 ($3.826\times10^{-1}$) | 0.121 ($3.107\times10^{-1}$) |
| **Distance OE1 E152-Mg2+ (Chain A)** | 0.021 ($8.629\times10^{-1}$) | -0.131 ($2.733\times10^{-1}$) |
| **Distance OE1 E152-Mg2+ (Chain C)** | 0.234 ($4.835\times10^{-2}$) | -0.066 ($5.811\times10^{-1}$) |
| **Distance OD2 D116-Mg2+ (Chain A)** | 0.271 ($2.107\times10^{-2}$) | -0.524 ($2.306\times10^{-6}$) |
| **Distance OD2 D116-Mg2+ (Chain C)** | 0.211 ($7.574\times10^{-2}$) | -0.423 ($2.103\times10^{-4}$) |

**Table S2. Spearman correlation coefficients between folding stability or binding energy and structural descriptors.** Structural descriptors include distances between the oxygen atoms of catalytic triad and $Mg^{2+}$ ions, as well as the RMSD of the 140–149 loop. Corresponding *p*-values are reported in parentheses, and statistically significant correlations (p-value < 0.01) are highlighted in red.

| Structural descriptor | Spearman correlation with folding stability (p-value) | Spearman correlation with binding energy (p-value) |
|---|---|---|
| **RMSD chain A** | 0.217 ($6.691 \times 10^{-2}$) | -0.058 ($6.308 \times 10^{-1}$) |
| **RMSD chain C** | 0.077 ($5.230 \times 10^{-1}$) | 0.141 ($2.390 \times 10^{-1}$) |
| **Distance OE1 E152-Mg2+ (Chain A)** | 0.038 ($7.488 \times 10^{-1}$) | -0.176 ($1.396 \times 10^{-1}$) |
| **Distance OE1 E152-Mg2+ (Chain C)** | 0.027 ($8.233 \times 10^{-1}$) | 0.224 ($5.903 \times 10^{-2}$) |
| **Distance OD2 D116-Mg2+ (Chain A)** | 0.208 ($7.986 \times 10^{-2}$) | -0.450 ($7.273 \times 10^{-6}$) |
| **Distance OD2 D116-Mg2+ (Chain C)** | 0.201 ($8.981 \times 10^{-2}$) | -0.431 ($1.588 \times 10^{-4}$) |

### Supplementary Figures

**Figure S1. Effect of individual drug-resistance mutations (DRMs) on folding stability and DNA binding, and evaluation of epistatic effects.** (A) Percentage of DRM variants showing increased or decreased stability compared to the WT, calculated relative to the total number of variants carrying the specific DRM. (B) Percentage of DRM variants showing increased or decreased DNA binding compared to the WT, calculated relative to the total number of variants carrying the specific DRM. (C) Percentage of DRM variants showing increased or decreased stability and DNA binding compared to the WT, calculated relative to the total number of variants carrying those specific DRMs.

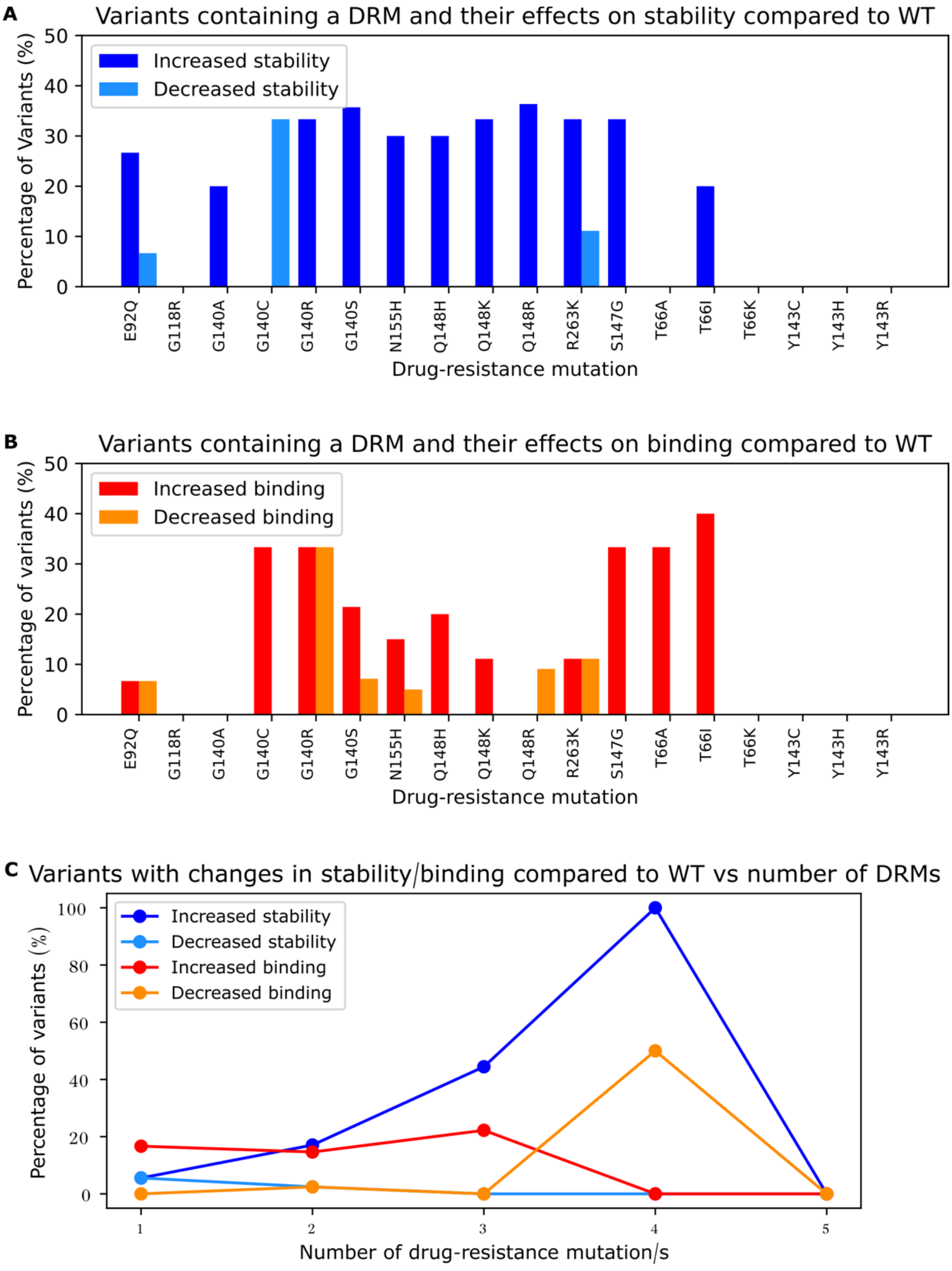

**Figure S2. Distances between oxygen atoms in the catalytic triad and $Mg^{2+}$ ions for chain A of HIV-1 integrase variants with and without drug-resistance mutations**. Each plot shows the distance between a specific oxygen atom and $Mg^{2+}$. On the left of each plot, a schematic representation shows the atoms considered for distance calculation. Each variant, including the WT, was analyzed in five independent replicates, and the plot shows the mean values and standard errors (error bars). Variants discussed in greater detail in the main text are highlighted with colored bars.

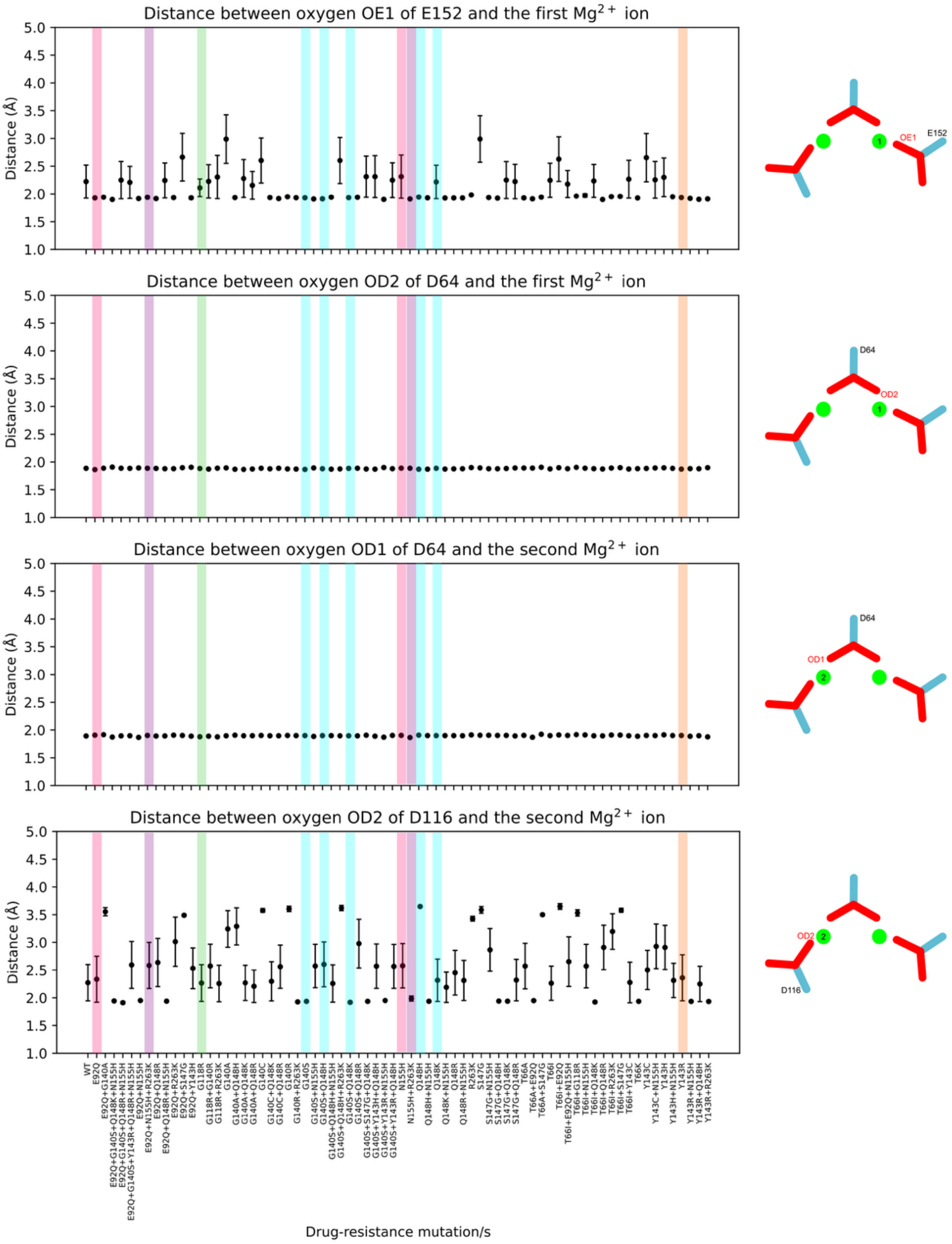

**Figure S3. Distances between oxygen atoms in the catalytic triad and $Mg^{2+}$ ions for chain C of HIV-1 integrase variants with and without drug-resistance mutations**. Each plot shows the distance between a specific oxygen atom and $Mg^{2+}$. On the left of each plot, a schematic representation shows the atoms considered for distance calculation. Each variant, including the WT, was analyzed in five independent replicates, and the plot shows the mean values and standard errors (error bars). Variants discussed in greater detail in the main text are highlighted with colored bars.

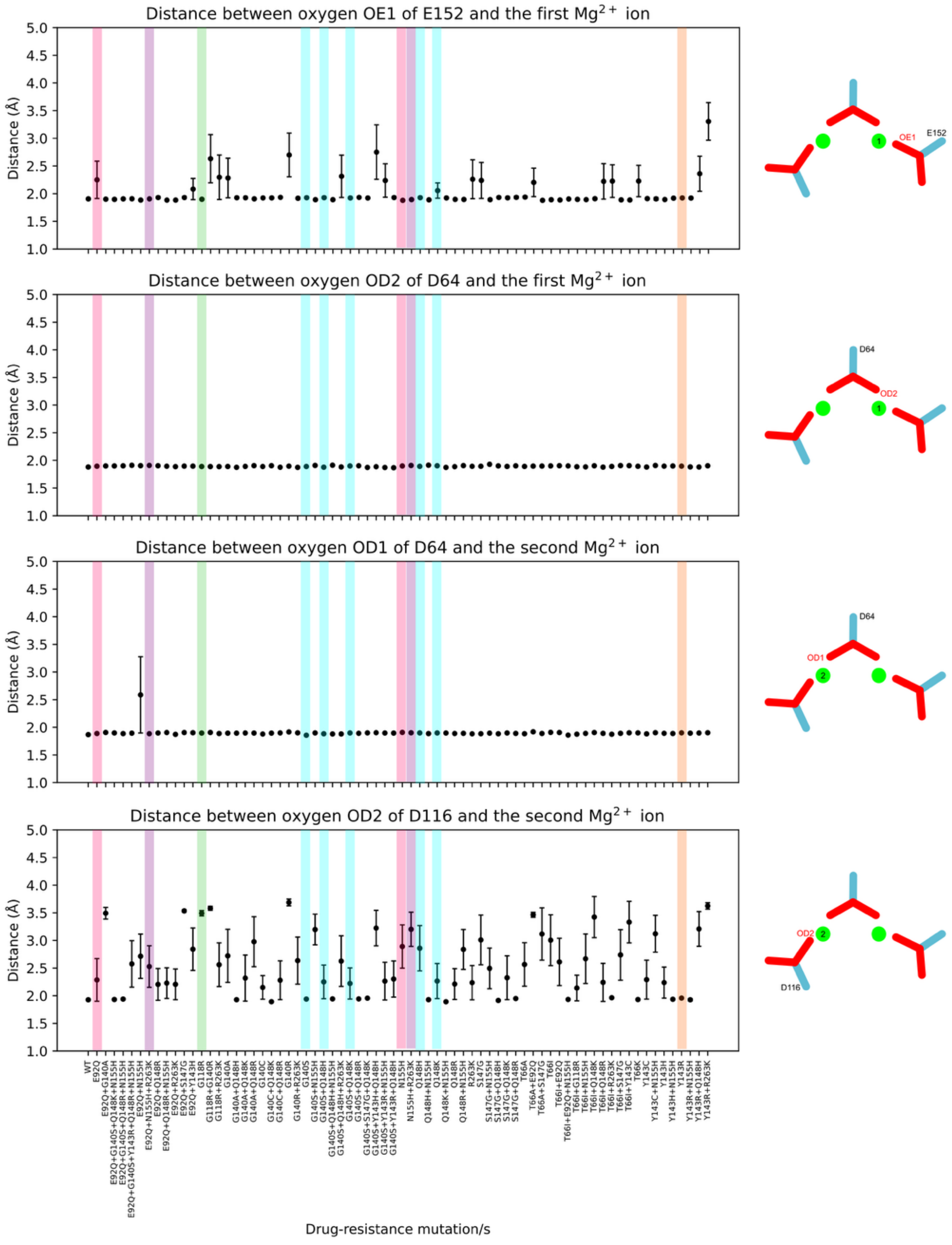

**Figure S4. Root mean squared deviation of loop 140-149 in chains A and C.** The plots show the root mean squared deviation (RMSD) of the loop 140-149 for chains A (above) and C (below). For each variant, the structure at the last step of energy minimization was used as the reference structure for the RMSD calculation. Each variant, including the WT, was analyzed in five independent replicates, and the plots show the mean values and standard errors (error bars).

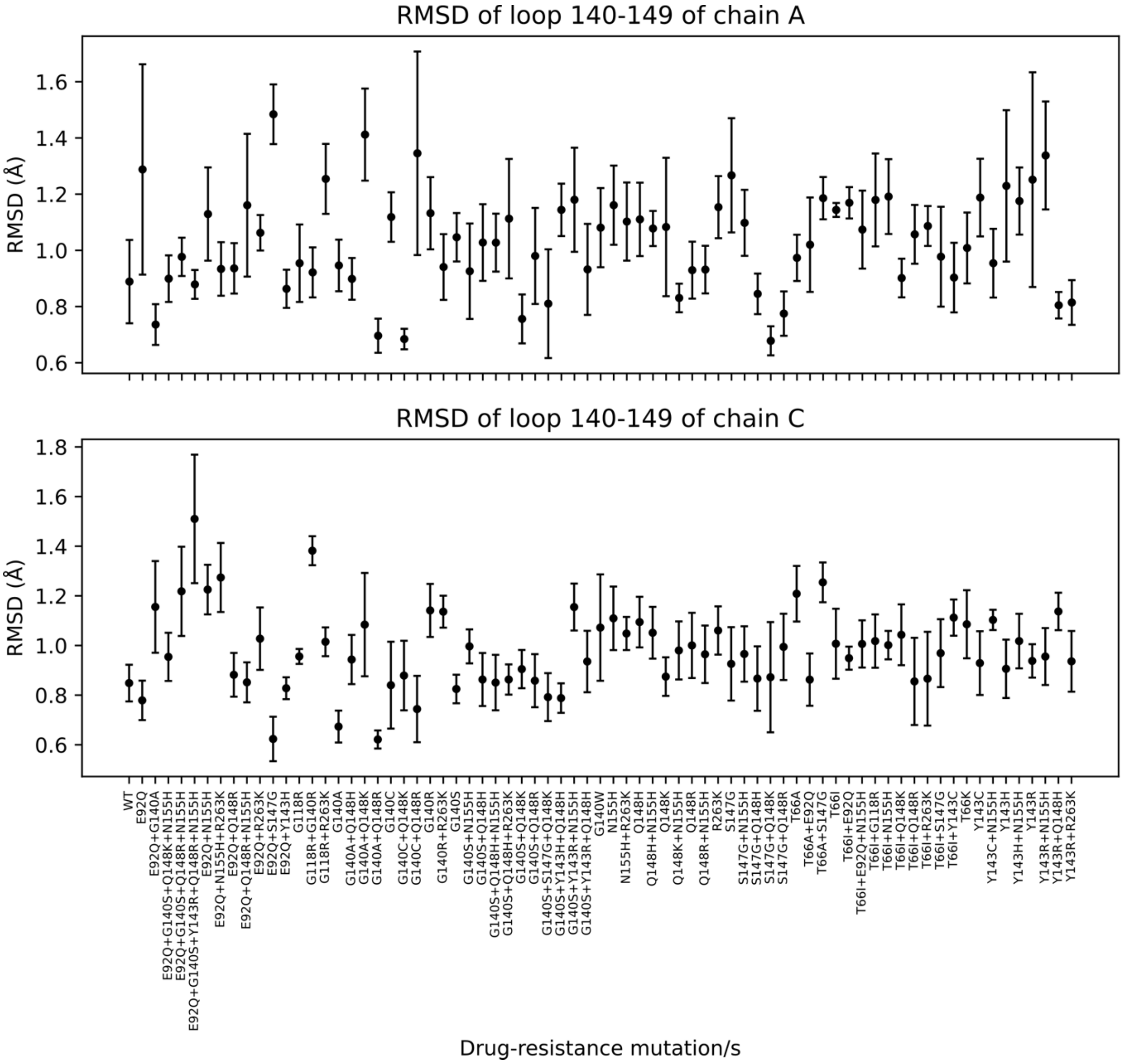

**Figure S5. Variation in folding stability of extant and reconstructed ancestral HIV-1 integrase variants relative to their immediate ancestor.** Phylogenetic tree illustrating the change in folding stability of extant (tip nodes) and reconstructed ancestral (internal nodes) HIV-1 IN variants relative to their immediate ancestor. Each tip node is labeled with the corresponding accession code from the Stanford HIV Drug Resistance Database (HIVdb), followed by the annotated HIV-1 group and subtype (e.g., AF457065_MA refers to HIVdb code AF457065, HIV-1 group M and, subtype A). An HIV-2 IN variant was included as an outgroup to root the tree. Phylogenetic branches are colored according to the variation of folding stability of each variant relative to its immediate ancestor, as indicated in the legend. Note that higher ΔG (lower stability), similar ΔG, and lower ΔG (higher stability) than the immediate ancestor are indicated in red, magenta, and blue, respectively.

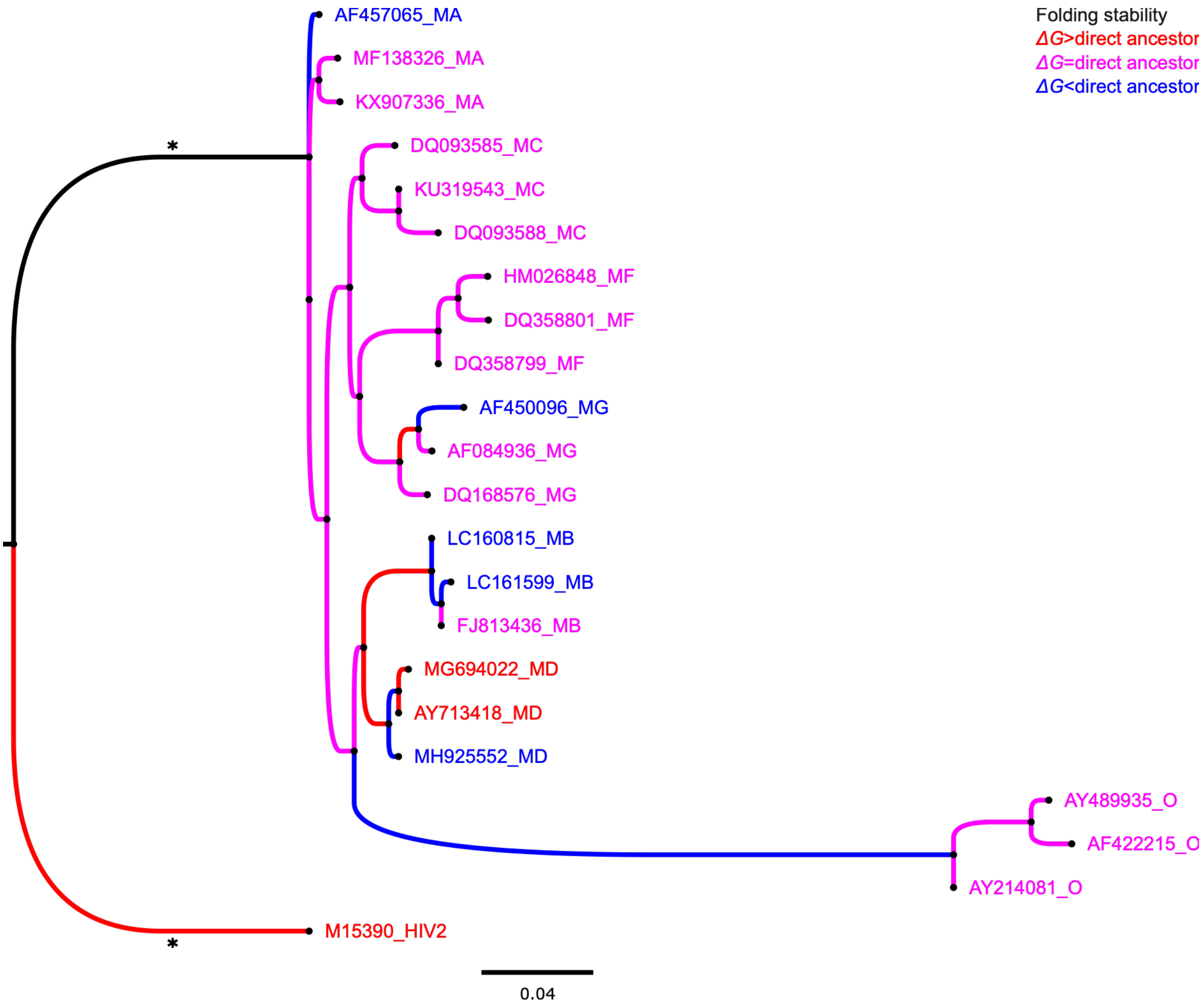

**Figure S6. Variation in binding free energy of extant and reconstructed ancestral HIV-1 integrase variants relative to their immediate ancestor.** Phylogenetic tree illustrating the change in binding free energy of extant (tip nodes) and reconstructed ancestral (internal nodes) HIV-1 IN variants relative to their immediate ancestor. Each tip node is labeled with the corresponding accession code from the Stanford HIV Drug Resistance Database (HIVdb), followed by the annotated HIV-1 group and subtype (e.g., AF457065_MA refers to HIVdb code AF457065, HIV-1 group M and, subtype A). An HIV-2 IN variant was included as an outgroup to root the tree. Phylogenetic branches are colored according to the variation of binding free energy of each variant relative to its immediate ancestor, as indicated in the legend. Note that higher ΔG (lower binding), similar ΔG, and lower ΔG (higher binding) than the immediate ancestor are indicated in red, magenta, and blue, respectively.

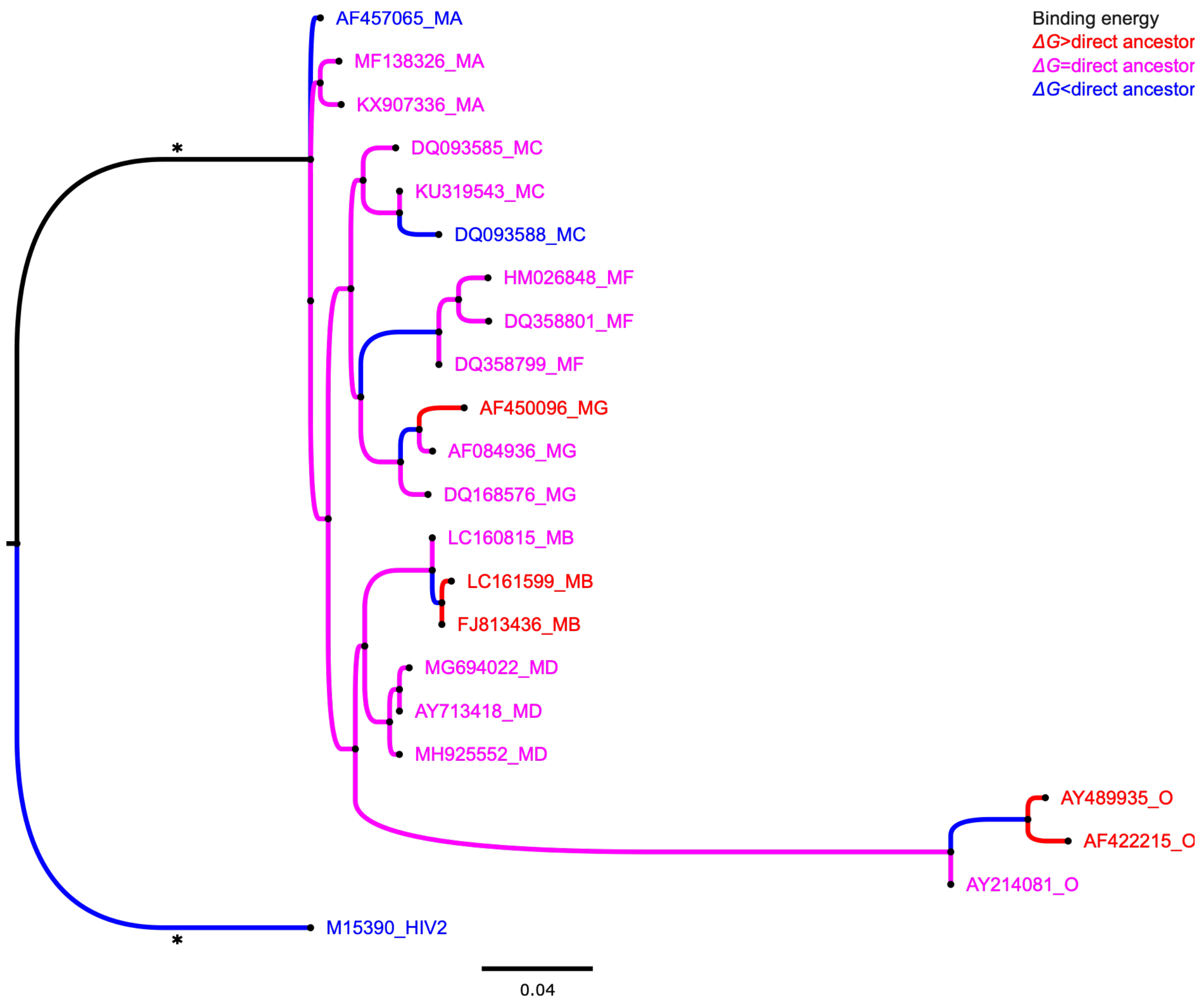

**Figure S7. Contribution of individual energy components to folding stability estimated with *FoldX*.** The plots show the average values of the different energetic terms (van der Waals, electrostatics, hydrophobic solvation, polar solvation, and van der Waals clash penalty) for each HIV-1 IN variant, calculated with *FoldX* from five independent replicates. Error bars indicate the corresponding standard error.

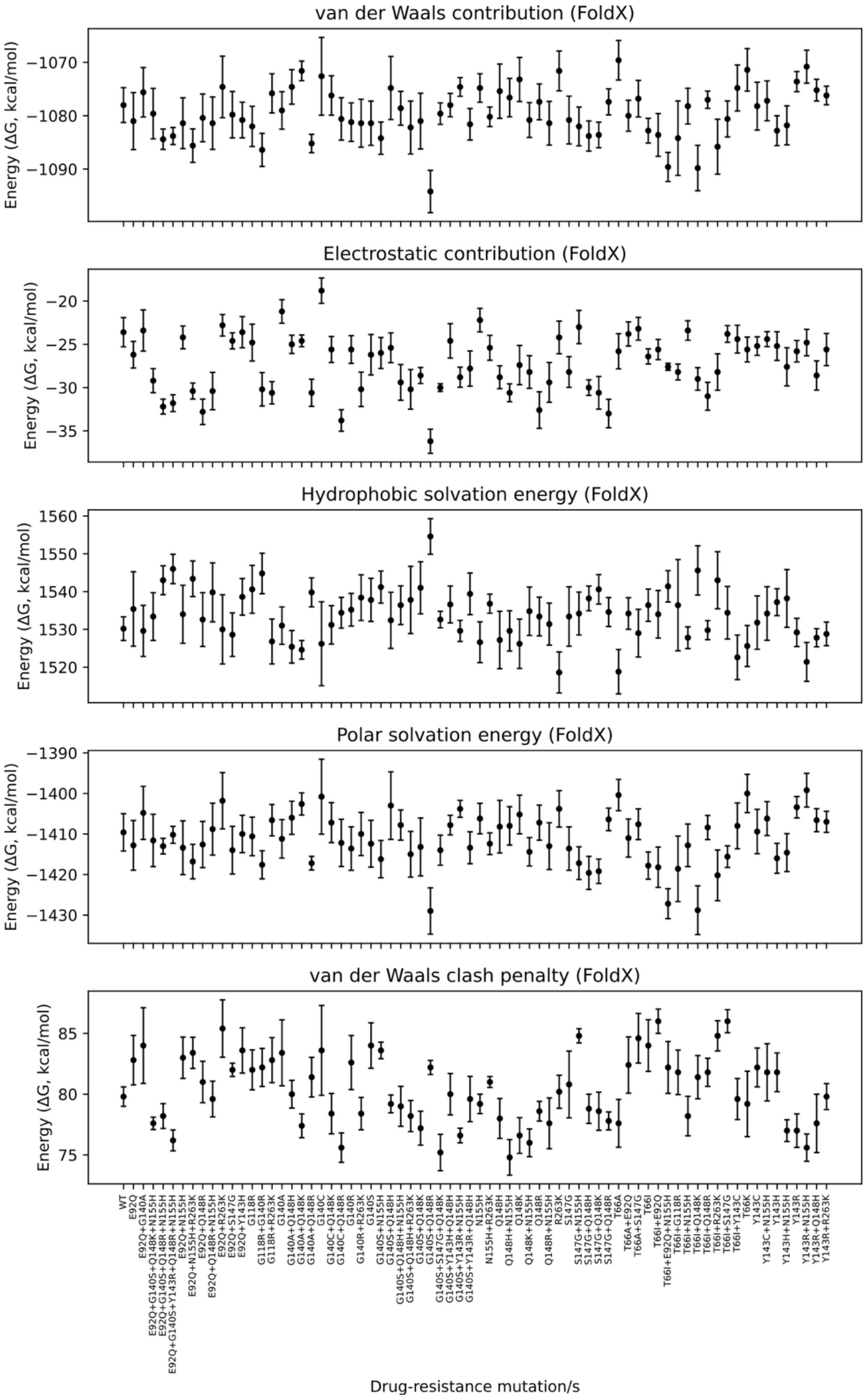

**Figure S8. Contribution of individual energy components to folding stability estimated with MMGBSA calculations.** The plots show the average values of the different energetic terms (van der Waals, electrostatics, hydrophobic and polar solvation) for each HIV-1 IN variant, calculated from five independent replicates. Error bars indicate the corresponding standard error.

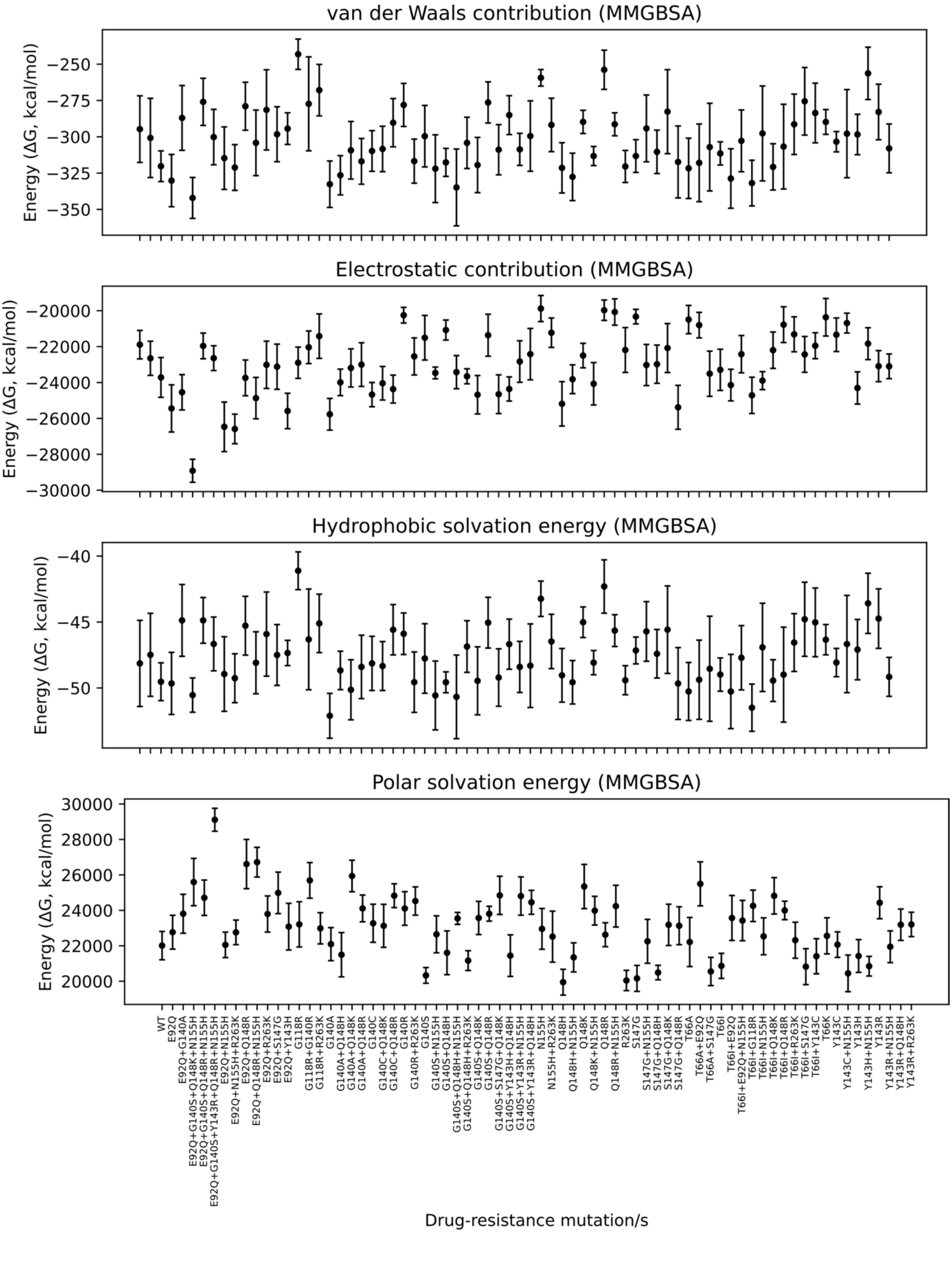
