## Supplementary Information 2 for "The evolution of the HIV-1 integrase through constraints from folding stability and functional interactions"

The supplementary material includes the Figures S2.1-S2.12 with analysis from molecular dynamics simulations.

**Figure S2.1.** Time evolution of the root-mean-square deviation (RMSD) for five independent replicas of each simulated system for analyzed DRMs.

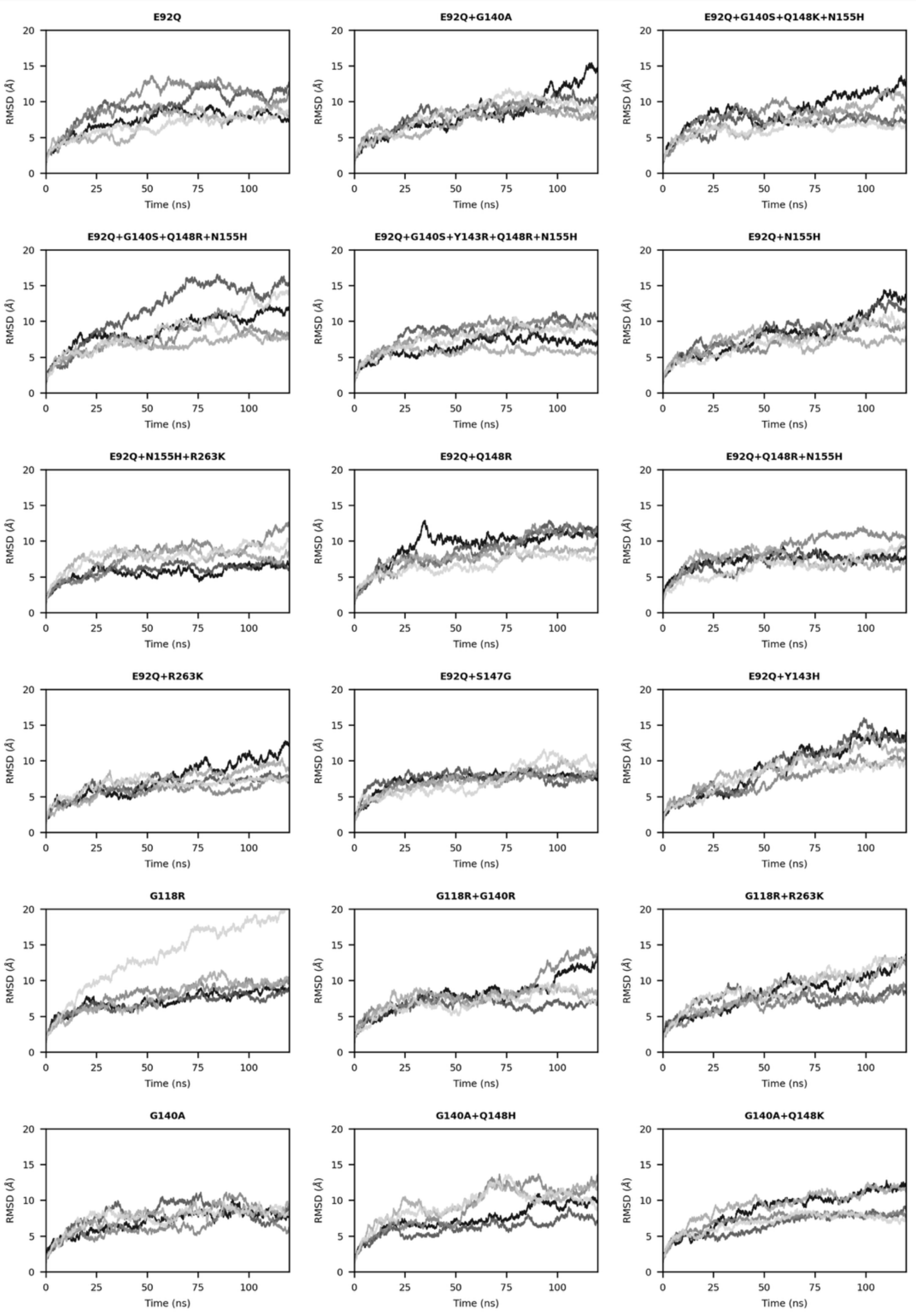

**Figure S2.2.** Time evolution of the root-mean-square deviation (RMSD) for five independent replicas of each simulated system for analyzed DRMs.

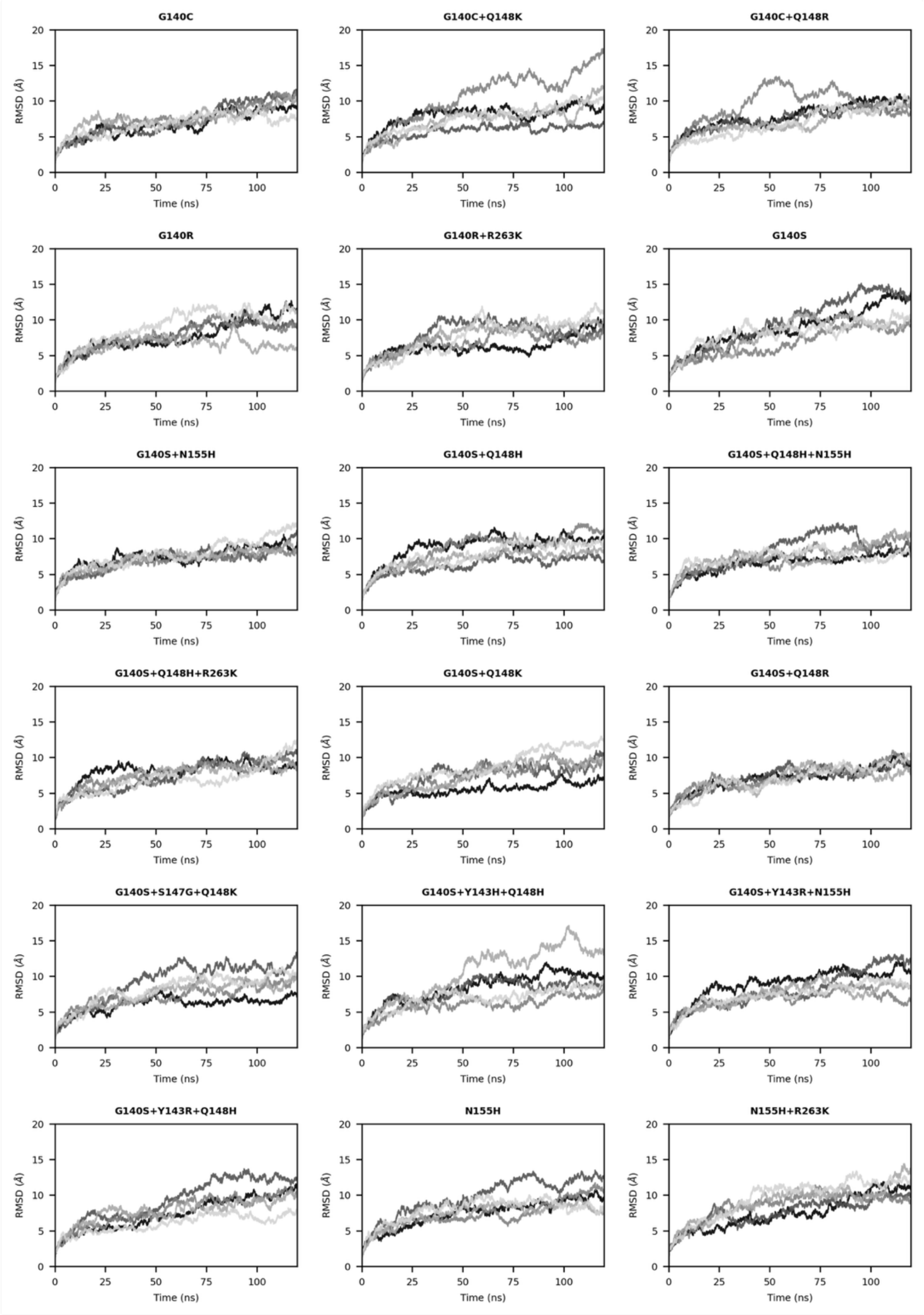

**Figure S2.3.** Time evolution of the root-mean-square deviation (RMSD) for five independent replicas of each simulated system for analyzed DRMs and the wild type (WT).

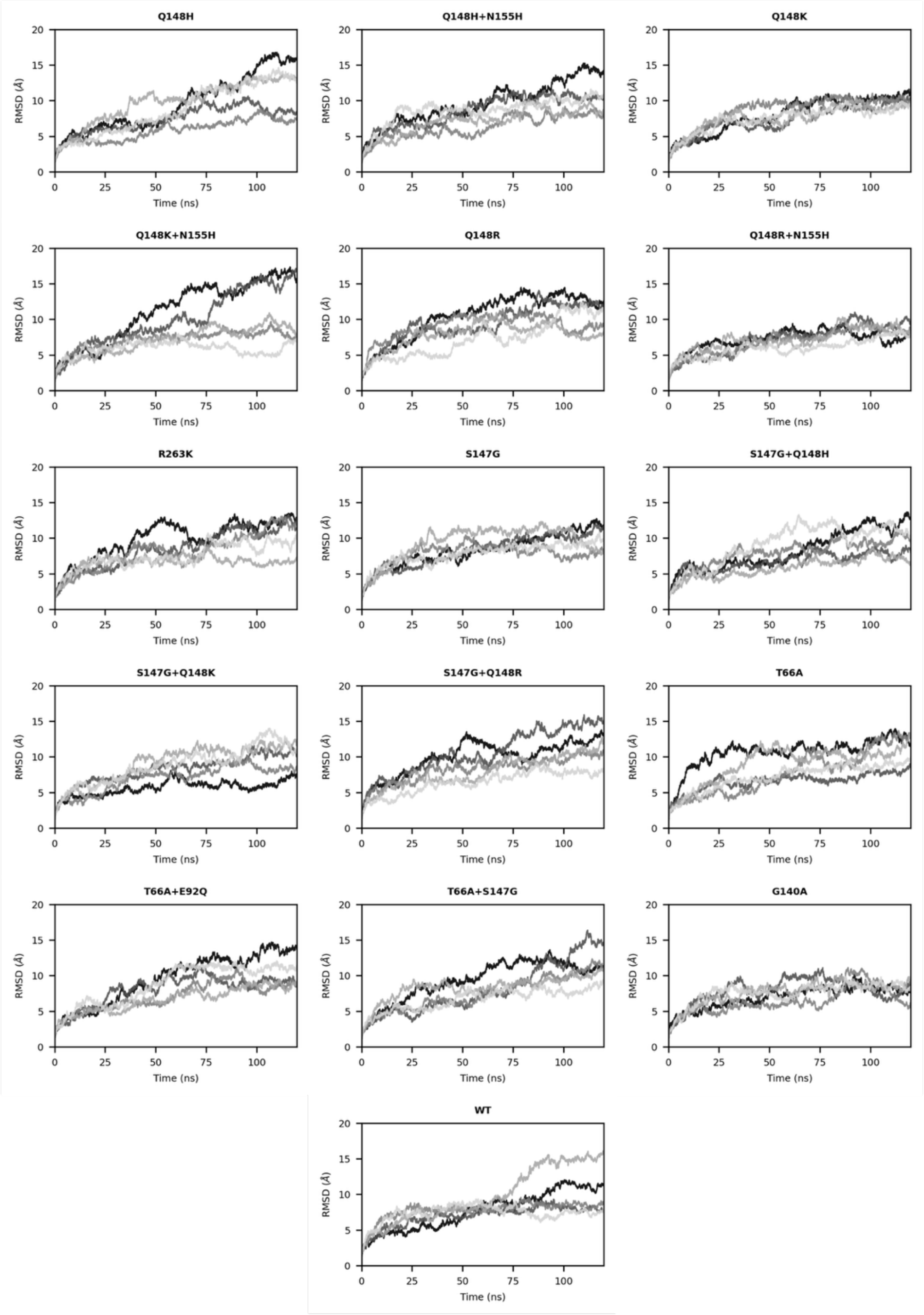

**Figure S2.4.** Time evolution of the root-mean-square deviation (RMSD) for five independent replicas of each simulated system for analyzed DRMs.

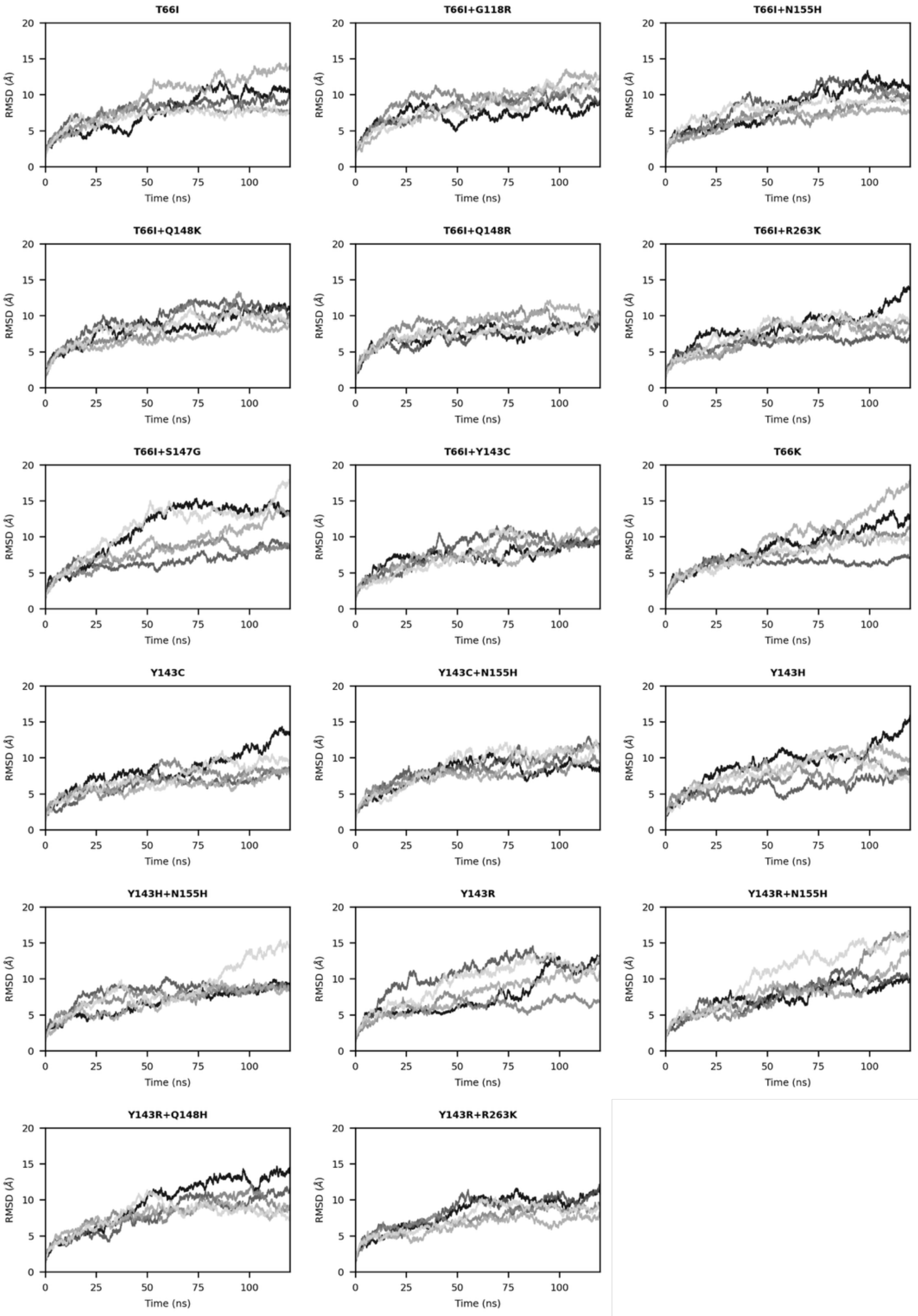

**Figure S2.5.** Time evolution of the root-mean-square deviation (RMSD) for five independent replicas of each simulated system for ancestral reconstructed sequences and the HIV-2 variant.

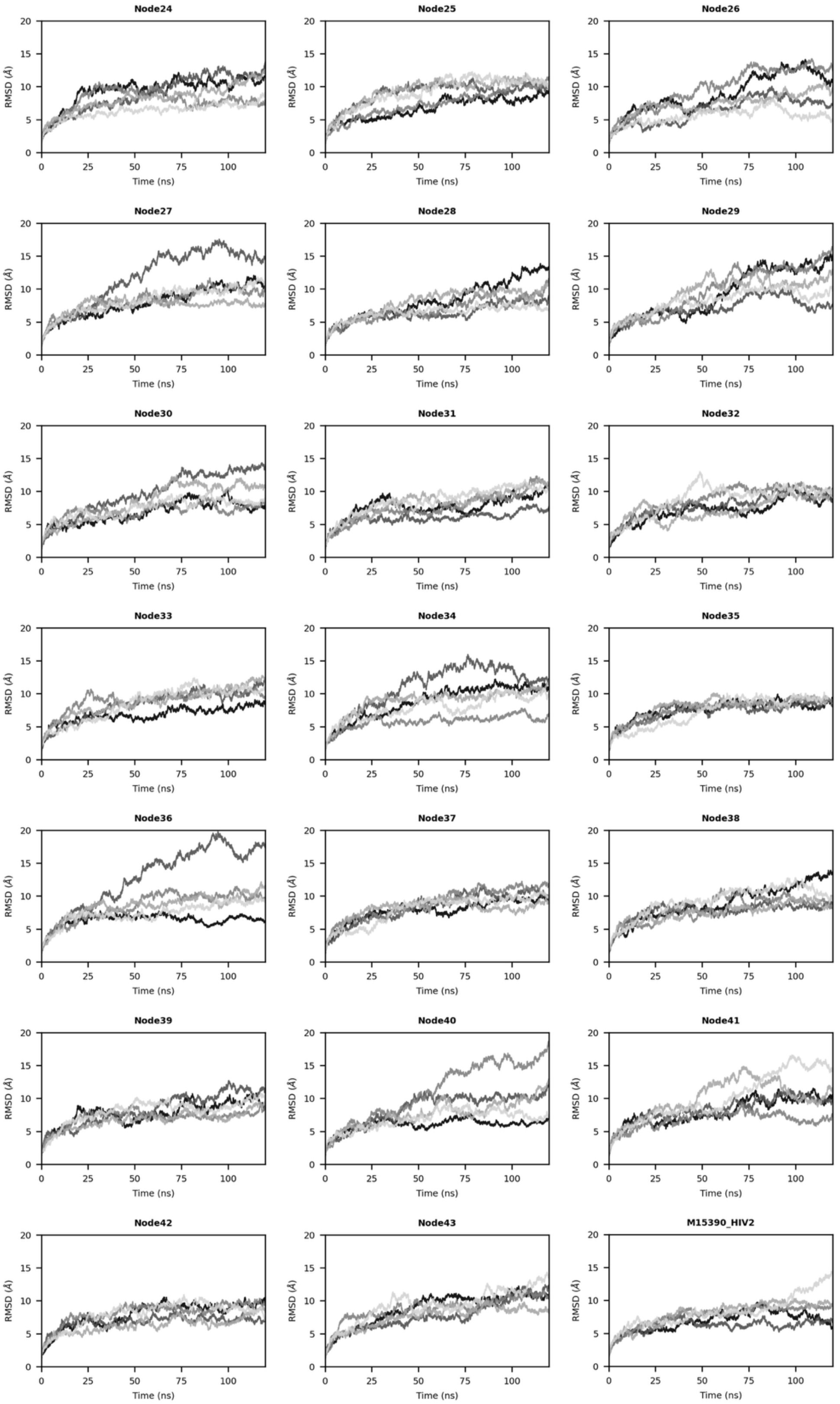

**Figure S2.6.** Time evolution of the root-mean-square deviation (RMSD) for five independent replicas of each simulated system for extant sequences.

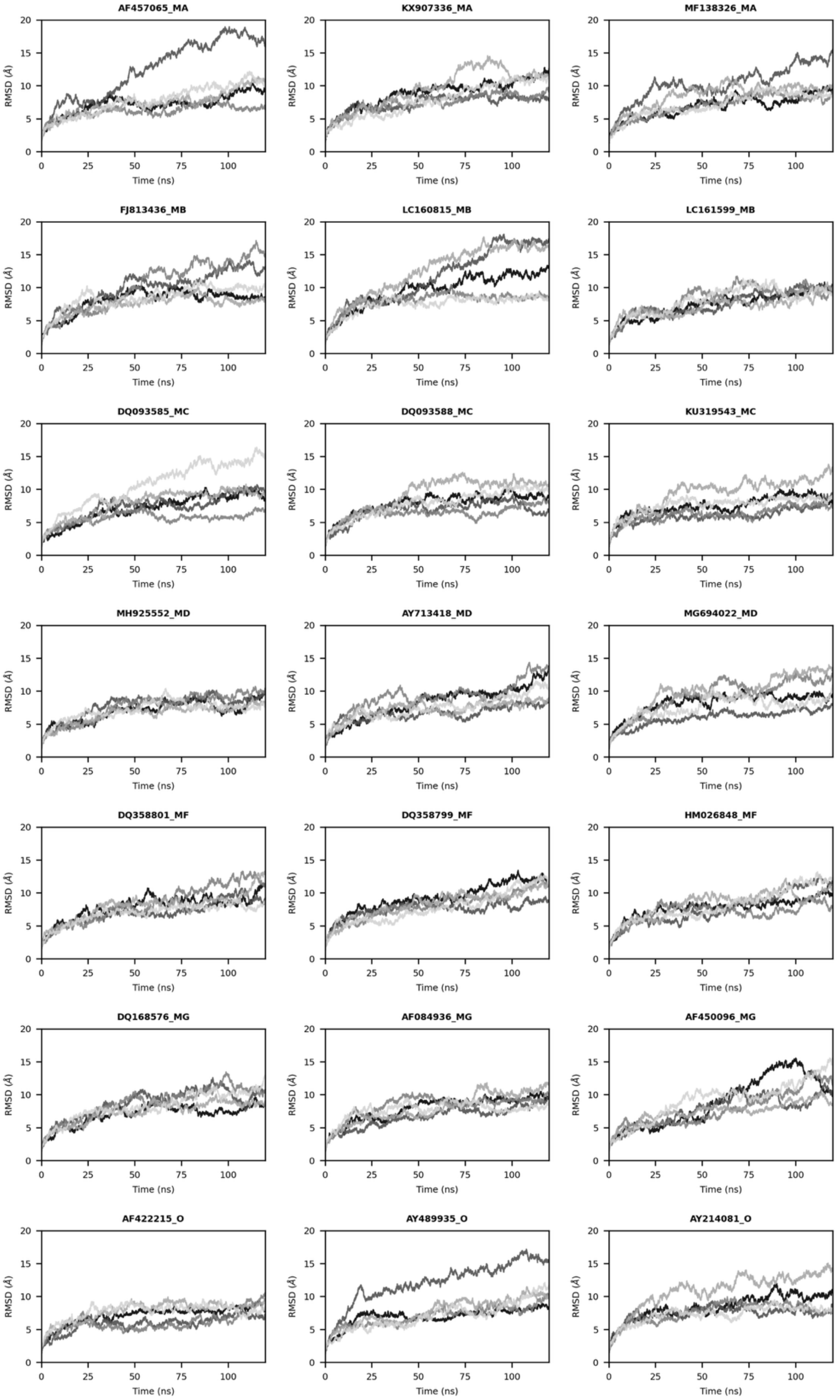

**Figure S2.7.** Root mean square fluctuations (RMSF) for the analyzed drug-resistant mutations (DRMs). RMSF values were averaged across the four monomers (270 residues each) and are reported as the mean (black line) and standard error (shaded gray area) across five independent replicas for each simulated system.

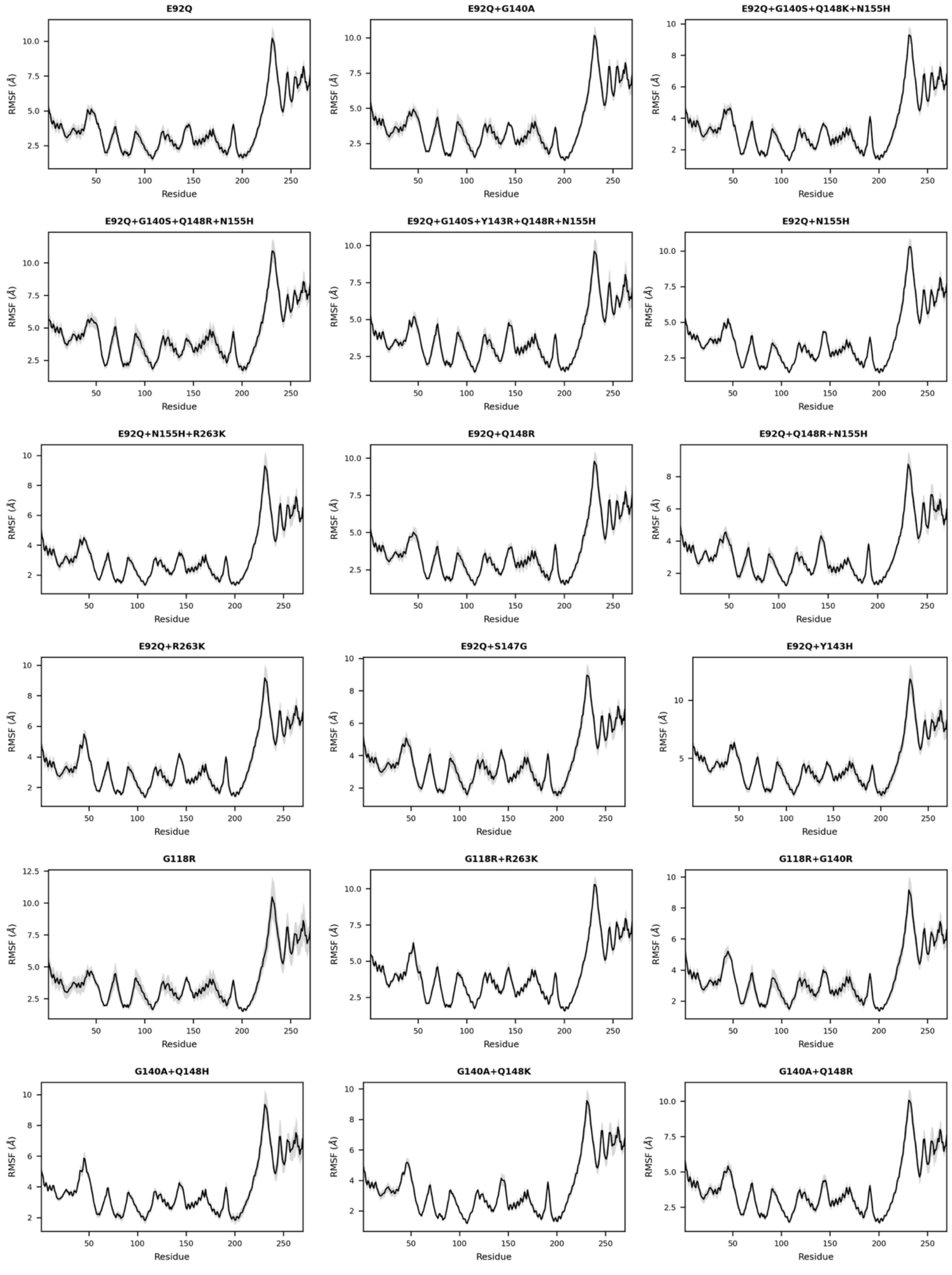

**Figure S2.8.** Root mean square fluctuations (RMSF) for the analyzed drug-resistant mutations (DRMs). RMSF values were averaged across the four monomers (270 residues each) and are reported as the mean (black line) and standard error (shaded gray area) across five independent replicas for each simulated system.

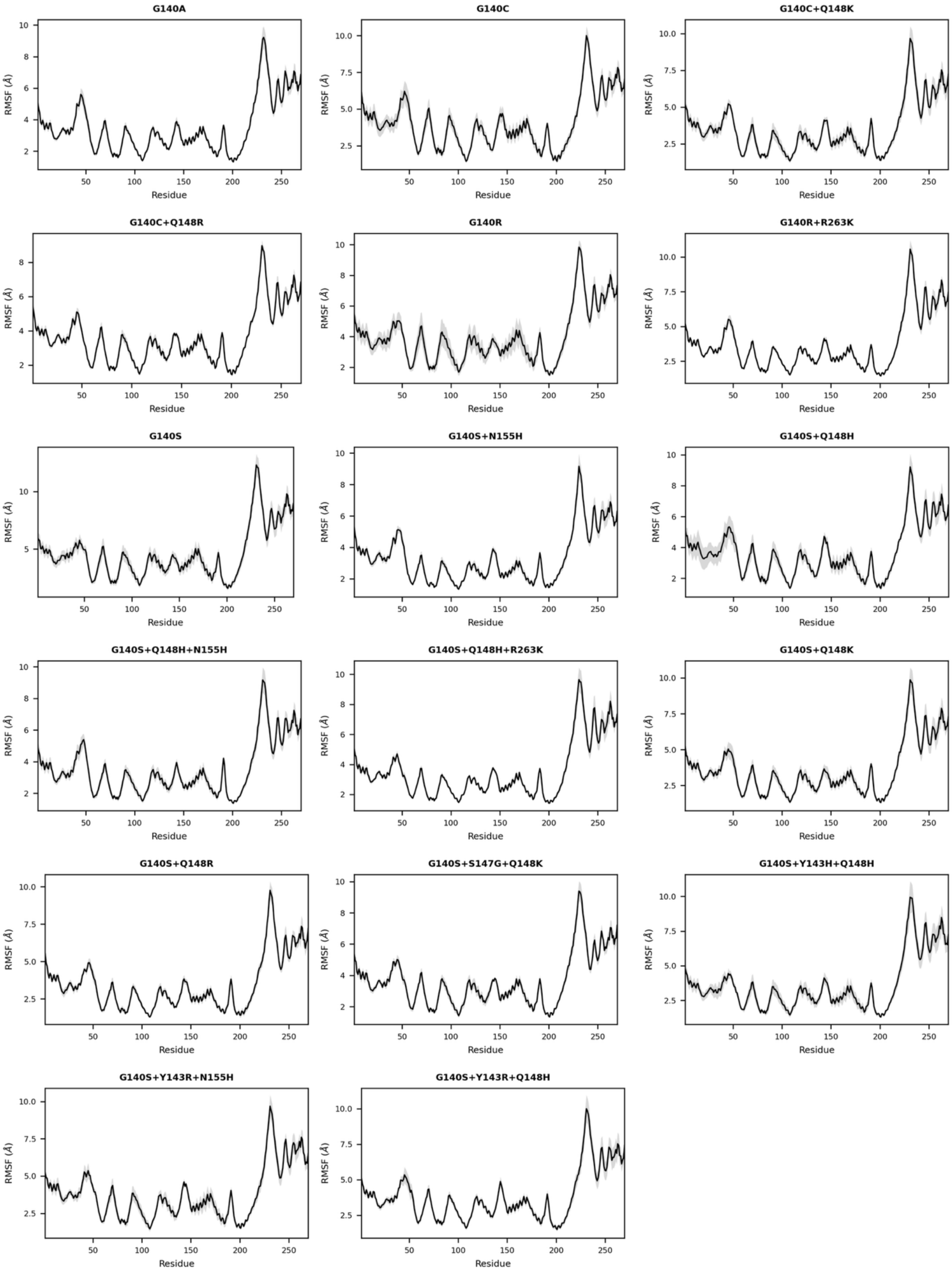

**Figure S2.9.** Root mean square fluctuations (RMSF) for the analyzed drug-resistant mutations (DRMs). RMSF values were averaged across the four monomers (270 residues each) and are reported as the mean (black line) and standard error (shaded gray area) across five independent replicas for each simulated system.

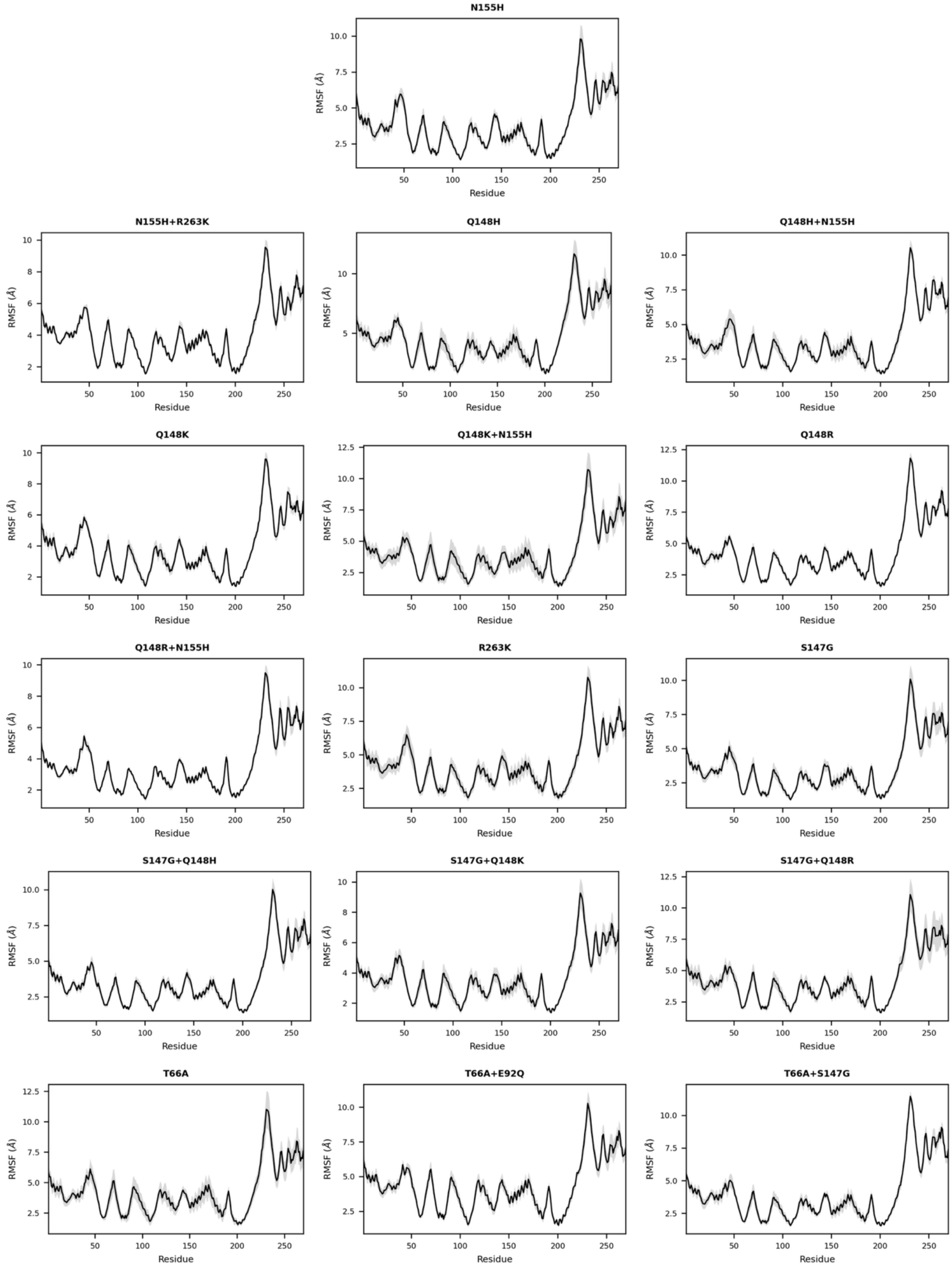

**Figure S2.10.** Root mean square fluctuations (RMSF) for the analyzed drug-resistant mutations (DRMs) and the wild type (WT). RMSF values were averaged across the four monomers (270 residues each) and are reported as the mean (black line) and standard error (shaded gray area) across five independent replicas for each simulated system.

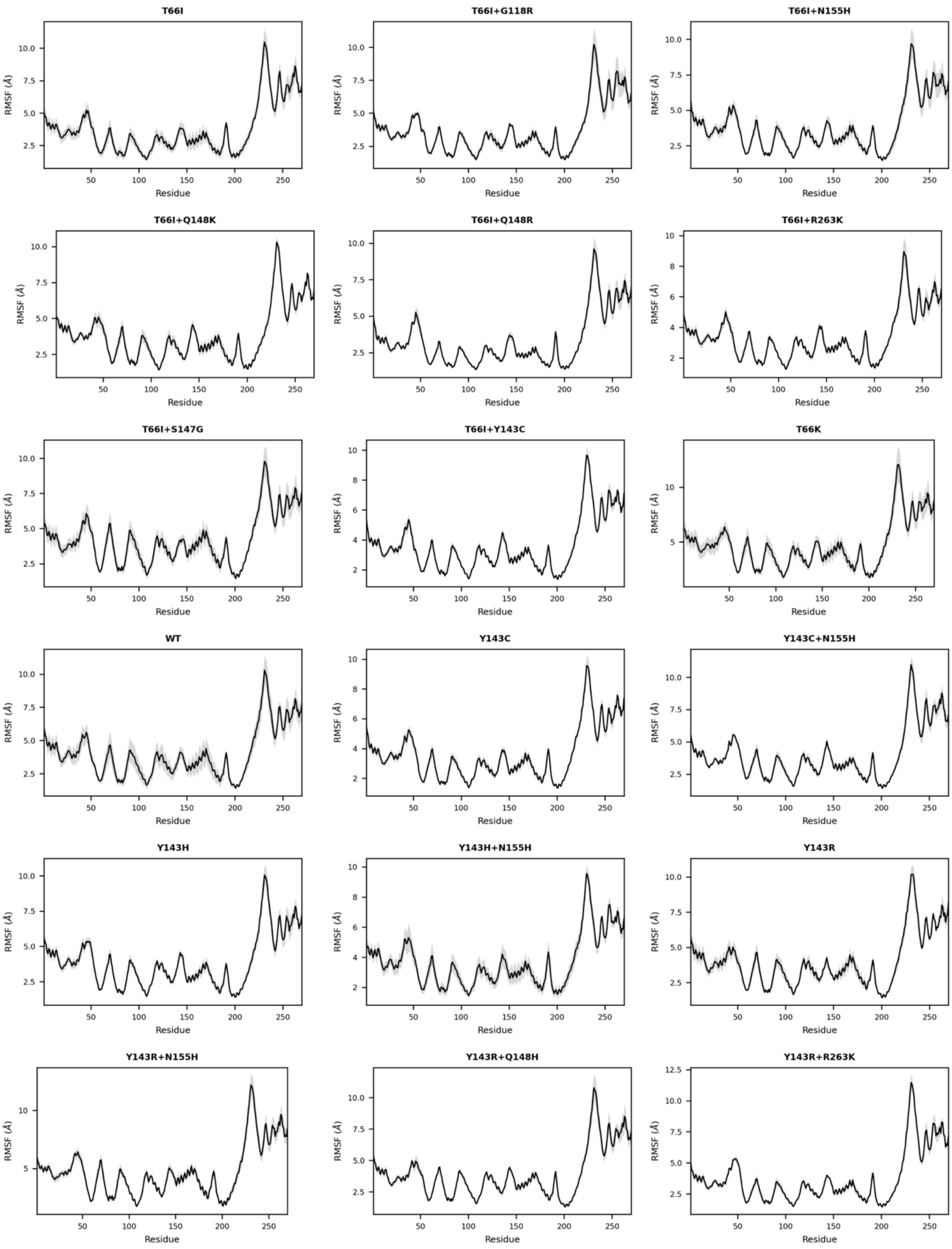

**Figure S2.11.** Root mean square fluctuations (RMSF) for ancestral reconstructed sequences and the HIV-2 variant. RMSF values were averaged across the four monomers (270 residues each) and are reported as the mean (black line) and standard error (shaded gray area) across five independent replicas for each simulated system.

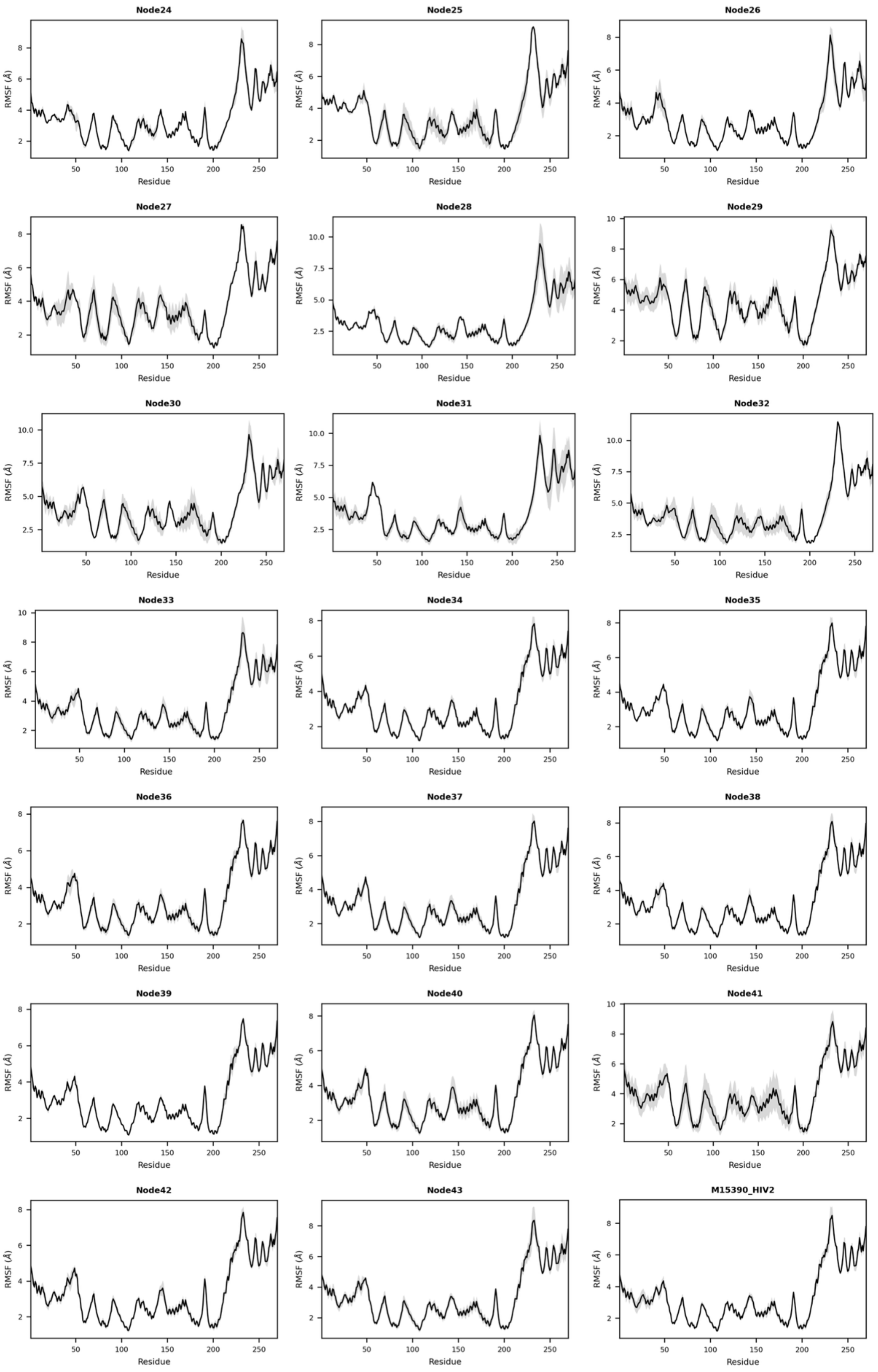

**Figure S2.12.** Root mean square fluctuations (RMSF) for extant sequences. RMSF values were averaged across the four monomers (270 residues each) and are reported as the mean (black line) and standard error (shaded gray area) across five independent replicas for each simulated system.

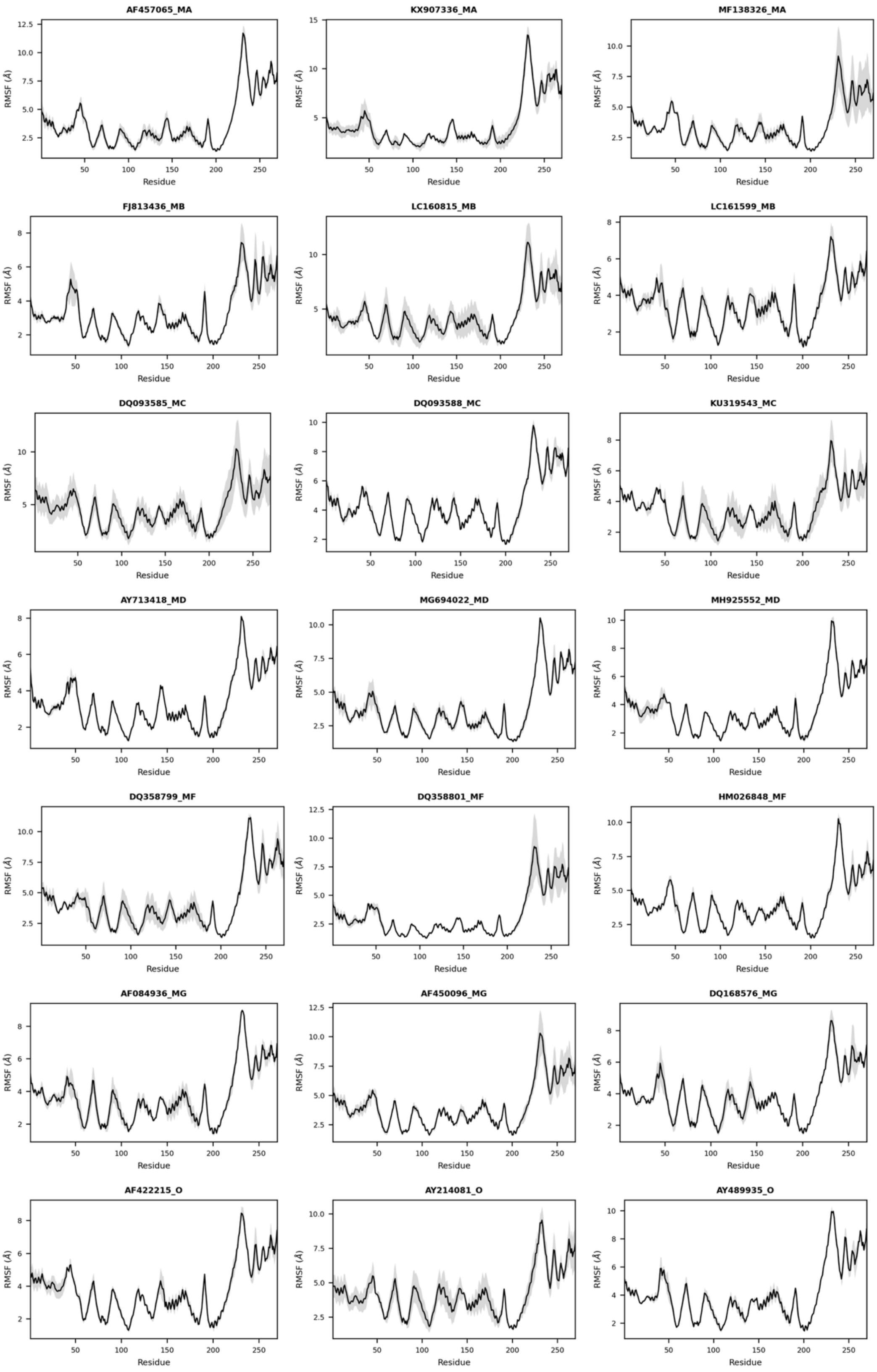
